# A multi-species toolkit of *TOP2* hypercleavage mutants for studying topoisomerase II–mediated DNA damage

**DOI:** 10.64898/2026.05.09.723998

**Authors:** David Ontoso, Monika Mehta, Aidin Shabro, John Dittmar, Robert J.D. Reid, Rodney Rothstein, John L. Nitiss, Scott Keeney

## Abstract

DNA topoisomerase II (TOP2) generates transient DNA double-strand breaks that are trapped as TOP2-DNA covalent complexes (TOP2cc) by antibiotic and chemotherapy drugs. Here, we characterize tools for study of cellular responses to TOP2cc, exploiting a *Saccharomyces cerevisiae TOP2* mutant (*TOP2-F1025Y,R1128G*) that generates spontaneous and inhibitor-induced covalent complexes at elevated frequencies. This Top2-hc (for “hypercleavage”) mutant protein inhibits yeast cell growth when expressed alone or with endogenous Top2, and growth defects are exacerbated in DNA-repair-deficient genetic backgrounds and/or in the presence of low doses of the Top2 poison mAMSA. We generated analogous mutations in human and mouse TOP2A and TOP2B that gave increased TOP2cc, hypersensitization to topoisomerase poisons, increased DNA damage, and decreased cell survival in cultured cells. We further established knock-in mouse models with inducible, tissue-specific expression of each TOP2-hc isoform, demonstrating overt organismal toxicity and cellular markers of DNA damage responses. To illustrate the potential of these genetic tools, we carried out proof-of-principle screens in yeast and cultured human cells for sensitivity to TOP2-hc. The yeast screen revealed strong requirements for homologous recombination, moderate roles for sister chromatid cohesion and kinetochore function, and dependencies on vesicle and vacuolar functions. The pilot shRNA screen in human cells revealed shared requirements for resistance to expression of either TOP2A-hc or TOP2B-hc as well as examples of isoform specificity. These findings establish hypercleavage mutant proteins as effective tools for studying topoisomerase isoform-specific DNA damage and offer a foundation for exploring TOP2cc toxicity and tolerance *in vivo*.

**Significance statement:** DNA topoisomerase II enzymes untangle chromosomes by cutting DNA, but incomplete resealing creates toxic damage that is the basis of antibacterial and chemotherapy drugs. Here we describe toolkits in yeast, mammalian cells, and mice that take advantage of mutant topoisomerase II enzymes that trap on DNA without drugs, creating powerful genetic systems to better study how cells deal with this type of DNA damage. We provide benchmarking data to validate these tools and to illustrate how they can be used for screens in cultured cells or tissue-specific experiments *in vivo*. These toolkits overcome longstanding technical barriers and enable new ways to study topoisomerase II-mediated DNA damage.

## Introduction

Type II topoisomerases generate reversible DNA double-strand breaks (DSBs) that resolve topological constraints inherent to replication, transcription, and chromosome segregation (1). DNA strand breakage occurs through a transesterification reaction in which a tyrosine residue carries out a nucleophilic attack on the DNA backbone, simultaneously severing the DNA and forming a tyrosyl phosphodiester link to the 5′ DNA end. Delay or failure to religate the transient TOP2-DNA covalent complex (TOP2cc) may convert this physiological intermediate into a genotoxic lesion (2). This intrinsic risk is exploited by cytotoxic drugs, known as topoisomerase poisons, that trap TOP2cc by shifting the cleavage–religation equilibrium (3). Widely used type II topoisomerase poisons include fluoroquinolone antibiotics and anticancer agents such as etoposide, doxorubicin and mitoxantrone (3).

TOP2cc lesions are particularly challenging because the DSB retains a bulky protein adduct covalently attached to the DNA end, thereby shielding the phosphotyrosyl bond from direct enzymatic access (4). Repair proceeds through partially redundant pathways, including nucleolytic processing by the MRE11–RAD50–NBS1 (MRN) complex and CtIP, or direct reversal by tyrosyl-DNA phosphodiesterase 2 (TDP2), frequently following proteolytic debulking (4, 5). Following TOP2 removal, the resulting DSBs are repaired via homologous recombination or nonhomologous end joining (NHEJ), depending on the DNA substrate, cell cycle stage, and organism (5, 6). Although many pathway components are defined, their relative contributions, redundancies, crosstalks and isoform-specific contributions remain incompletely understood.

Genetic analysis of sensitivity to topoisomerase poisons has been a fruitful approach (7–9), but some of these drugs cause toxicity through topoisomerase-independent mechanisms, such as the generation of reactive oxygen species (10). Moreover, sensitivity may be strongly dependent on cell permeability and/or drug efflux (11). Additionally, mammalian cells express two paralogs of topoisomerase II — alpha (TOP2A) and beta (TOP2B) — which are mainly sensitive to the same poisons (5) but are also differentially targeted by some drugs, affecting their respective contributions to toxicity (10). Selective siRNA depletion has been used to study isoform-specific contributions to conversion of poison-induced TOP2cc into DSBs (12), but this strategy alters overall enzyme abundance and perturbs ongoing replication or transcription programs, making it challenging to disentangle isoform-intrinsic DNA cleavage behavior from secondary cellular responses.

An alternative strategy is the use of self-trapping topoisomerase mutants that behave as if constitutively poisoned, even in the absence of a drug. The original examples of such mutants are the *S. cerevisiae top1-103* (R420K) and *top1-T722A* alleles, which mimic the cytotoxic action of the type I topoisomerase poison camptothecin (13, 14). Such alleles have been used in yeast and mammalian cells to screen for genes required for resistance to TOP1cc, to study effects of TOP1cc on neuronal cells in culture, or as a tool to identify new poisons and inhibitors (15–18).

Analogously, *S. cerevisiae TOP2-F1025Y,R1128G* exhibits behaviors resembling constitutive poisoning by type II topoisomerase inhibitors (19). Here we extend characterization of this *TOP2-hc* mutation *in vivo* in yeast and construct analogous mutants of TOP2A and TOP2B in human and mouse, enabling isoform-specific generation of TOP2 cleavage complexes independent of protein abundance. This work establishes toolkits and proof-of-principle experimental systems for study of the consequences of and cellular responses to topoisomerase II-mediated DNA damage in *S. cerevisiae*, cultured human and mouse cells, and specific mouse tissues *in vivo*.

## Results

### Toxicity in yeast from inducible expression of hypercleavage mutant Top2 protein

*S. cerevisiae TOP2-F1025Y,R1128G* was originally isolated because of its hypersensitivity to mAMSA and etoposide (19, 20). The mutant protein has relaxation activity comparable to wild type *in vitro* but is self-poisoning, i.e., frequently becomes trapped on DNA. When constitutively overexpressed from a low-copy plasmid *in vivo*, the mutant triggers a hyper-recombination and mutagenic phenotype and is synthetically lethal with absence of the homologous recombination protein Rad52 (19). It was proposed that the mutations cause persistent DSBs by destabilizing the critical dimer interface known as the C-terminal (C) gate (***SI Appendix*, Fig. S1*A***).

To build a yeast toolkit, we constructed low-copy *ARS, CEN* plasmids expressing wild-type or mutant *TOP2* with a C-terminal triple FLAG tag under the control of a copper-inducible promoter (*P_CUP1_*) (***SI Appendix*, Fig. S1*B*)**. The basal copper in yeast synthetic medium (0.25 µM) gave expression comparable to endogenous Top2, i.e., yielding an approximately two-fold increase in total Top2 (**Fig. 1*A* and *SI Appendix*, Fig. S1*C***). Addition of 100 µM CuSO_4_ rapidly induced ∼3–4 fold more protein (**Fig. 1*A* and *SI Appendix*, Fig. S1*C* and S1*D*)**.

**Fig. 1.**
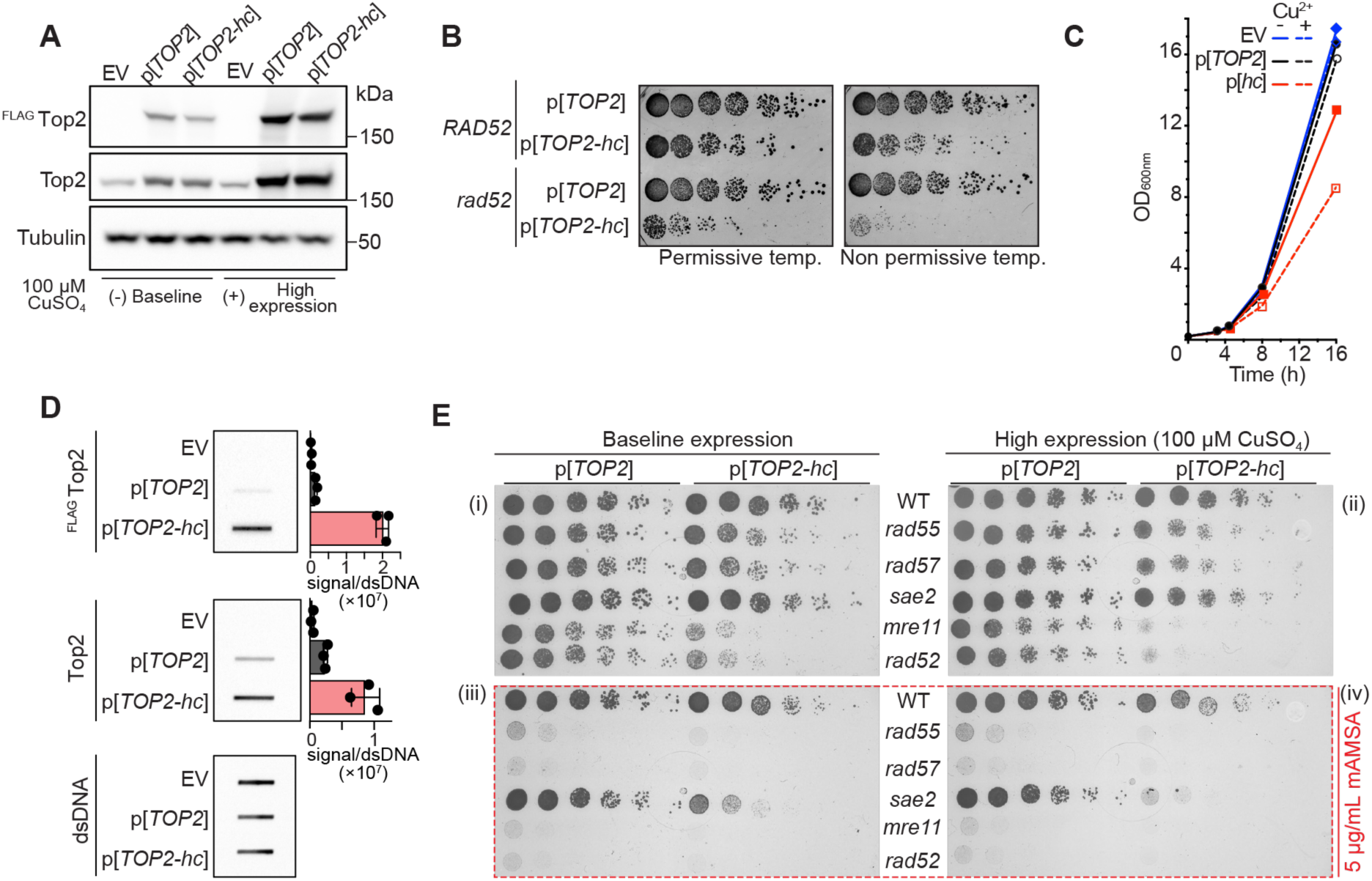
Phenotypic characterization of the self-poisoning *TOP2-hc* allele in yeast. (A) Immunoblot analysis of ^FLAG^ Top2 and ^FLAG^ Top2-hc expression. Baseline and induced (100 µM CuSO₄) expression levels are shown. Quantification is in ***SI Appendix*, Fig. S1*C***. (B) Compromised complementation of temperature-sensitive *top2-4* by *TOP2-hc* in a DNA repair-defective background. The indicated plasmids were transformed into *RAD52* or *rad52* deletion strains in a *top2-4* background. Fivefold serial dilutions were spotted onto selective medium plates and incubated at permissive (28 °C) or non-permissive (34 °C) temperatures. (C) Delayed growth rates with Top2-hc expression. Growth curves (optical density at 600 nm) are shown for a wild-type strain carrying the indicated plasmids cultured with or without supplementation with 100 µM CuSO_4_. (D) Increased Top2cc upon Top2-hc expression, detected by the ICE assay. Genomic DNA (0.5 µg per slot) was purified as described in (52), immobilized on a nitrocellulose membrane and immunodetected using anti-FLAG, polyclonal anti-Top2, and anti-double-stranded DNA. Quantification shows chemiluminescence signals normalized to dsDNA (mean ± SD from three independent experiments). (E) Hypersensitivity of DNA repair mutants to Top2-hc expression. Fivefold serial dilutions of exponentially growing cultures of the indicated strains were spotted on selective medium plates without (i, iii) or with (ii, iv) copper supplementation and without (i, ii) or with (iii, iv) mAMSA (5 µg/ml). Plates were scanned 42 hr after inoculation.

We introduced these plasmids into strains carrying the temperature-sensitive *top2-4* allele at the endogenous locus (21) and plated serial dilutions on copper-supplemented medium (**Fig. 1*B***). In a repair-proficient background (*RAD52+*), *TOP2-hc* overexpression decreased the number and size of colonies at a permissive temperature for *top2-4* (28 °C). At a non-permissive temperature (34 °C), growth was further suppressed, but the residual viability suggests that the mutant enzyme is sufficiently active to support viability as the sole source of topoisomerase II activity. Consistent with prior results (19), *TOP2-hc* overexpression gave a stronger growth defect at 28 °C in a repair-deficient background (*rad52*) and almost completely eliminated growth at 34 °C (**Fig. 1*B***).

Top2-hc also inhibited growth in a strain with wild-type endogenous *TOP2*, modestly with low baseline expression and to an even greater extent when overexpressed (**Fig. 1*C***). The hypercleavage phenotype is thus codominant with wild type, hence the upper-case designation for *TOP2-hc*. Top2-hc localized to nuclei, but its overexpression caused cellular morphologies associated with cytotoxicity (***SI Appendix* Fig. S1*E***). Top2-hc overexpression also led to increased Rad52-EGFP foci (***SI Appendix* Fig. S1*F***); activation of the DNA damage response kinase Rad53 (***SI Appendix* Fig. S1*G***); increased accumulation of modified Top2 species (likely sumoylation and/or ubiquitylation, *(**SI Appendix*** **Fig. S1*H***; and, consistent with prior results (19), higher levels of Top2cc in cells, as measured by the *in vivo* complex of enzyme (ICE) assay (**Fig. 1*D***).

Baseline Top2-hc expression compromised cell growth strongly in *mre11* and *rad52* backgrounds and moderately in *rad55* and *rad57* (Rad51 paralog mutants), but it unexpectedly had no effect in *sae2* (**Fig. 1*E*, panel i**). Top2-hc overexpression caused synthetic lethality in *mre11* and *rad52,* increased synthetic sickness in *rad55* and *rad57*, and mildly suppressed growth in *sae2* (**Fig. 1*E*, panel ii**). A low dose of mAMSA, tolerated by wild-type cells but lethal to all mutants except *sae2* (**Fig. 1*E*, panel iii**), resulted in slow growth in the *sae2* mutant when combined with baseline Top2-hc expression (**Fig. 1*E*, panel ii**) and was lethal with Top2-hc overexpression (**Fig. 1*E*, panel iv**).

Together with previous observations (19), these findings show that Top2-hc causes accumulation of Top2cc and associated toxicity in DSB repair-proficient cells without exogenous topoisomerase poisons. This toxicity is exacerbated by DNA repair deficiency or treatment with otherwise well-tolerated doses of topoisomerase poisons. Toxicity can be ameliorated by substoichiometric amounts of wild-type Top2. These studies establish a tunable system for creating Top2-mediated DNA damage *in vivo* in *S. cerevisiae*.

### Screening for sensitivity to Top2-hc in yeast

As proof of principle, we screened a yeast non-essential gene deletion collection for synthetic lethal interactions with *TOP2-hc*. An empty vector (control) or plasmids expressing either wild-type Top2 or Top2-hc were transferred into the deletion collection by selective ploidy ablation (SPA) (18). Each plasmid-bearing haploid deletion strain (n = 4565) was pinned in quadruplicate on synthetic complete plates lacking leucine (SC-LEU) to select for maintenance of the plasmid, with or without additional 100 µM CuSO_4_, and colony sizes were quantified. An overview is shown in ***SI Appendix* Fig. S2A, left** (full results in **Dataset S1**). Example arrayed plates are shown in ***SI Appendix*, Fig. S2*B***, with selected strains highlighted in ***SI Appendix*, Fig. S2*C***.

Based on these results, 523 ORF deletions were selected (absolute z-scores > 2) for a secondary validation screen in which 16 colonies were tested for each strain (***SI Appendix* Fig. S2*A*, middle, and Fig. S2*D*; Dataset S1**). Using the more robust output from the validation screen, 110 deletion mutants were hypersensitive to Top2-hc (log_2_ growth ratio < -0.5, p < 0.05) (**Fig. 2*A***, ***SI Appendix* Fig. S2*A*, right, S2*E*, S2*F*, and Dataset S1**). We also scored 28 suppressor mutants that surprisingly appeared to grow better in the presence of Top2-hc compared to the empty vector control (***SI Appendix* Fig. S2*A*, S2*E***), but these largely failed final targeted validation in spot tests (***SI Appendix*, Fig. S3**).

**Fig. 2.**
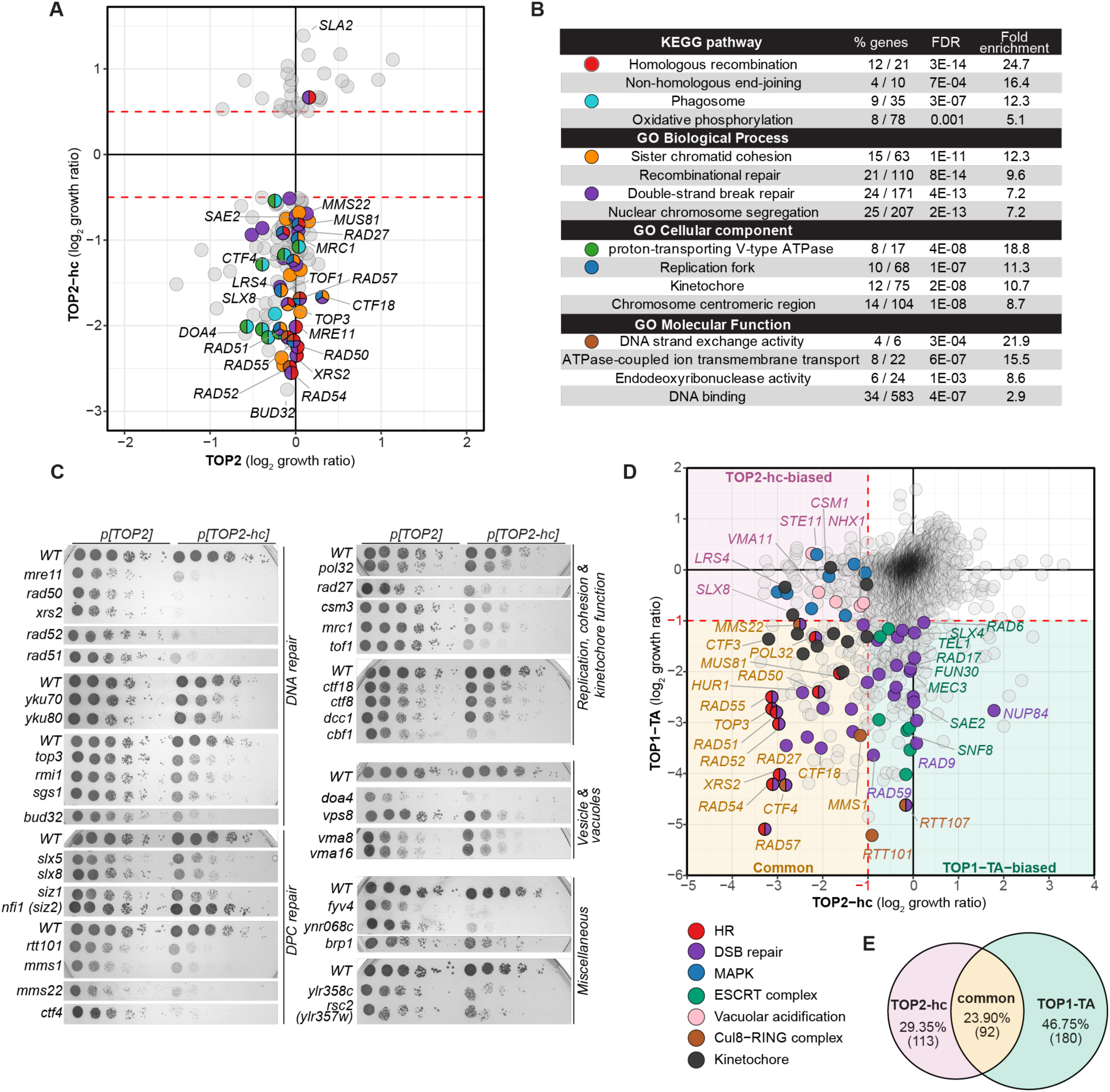
Genetic determinants of sensitivity to Top2-hc expression in yeast. (A) SPA screen results. The scatterplot shows log₂ growth ratios (EV:Top2 plasmid) under high expression conditions for Top2 (x axis) versus Top2-hc (y axis). Red dashed lines indicate log₂ growth ratio thresholds applied to the validation screen. Highlighted genes are colored according to functional categories in panel B. (B) Functional enrichment analysis of KEGG and Gene Ontology terms associated with Top2-hc sensitivity. FDR, false discovery rate. Full data are in **Dataset S2.** (C) Validation of Top2-hc hypersensitivity of selected mutants by spot-dilution assays. Fivefold serial dilutions of exponentially growing cultures were spotted on selective medium supplemented with copper to induce high expression. Mutants are grouped by function. (D) Comparison of Top2-hc and Top1-TA SPA screens. Quadrants are shaded to denote common sensitivity (lower left, orange), Top2-hc-biased (upper left, red) or Top1-TA-biased (lower right, green). Selected genes are color-coded by functional category. Full data are in Dataset S3. (E) Venn diagram summarizing overlap between Top2-hc and Top1-TA SPA screens (color coded as for quadrant shading in panel D).

As expected, copper-induced high expression further exacerbated cytotoxicity, increasing the cell growth defects in sensitive mutants (***SI Appendix*, Fig. S2*D* and S2*F***). For some mutants like *rad52* or *mre11*, even baseline expression without copper supplementation was enough to cause synthetic lethality (***SI Appendix*, Fig. S2*D* and S2*F***, see mutants on dashed diagonal line). High expression of wild-type Top2 had little effect on the viability of most strains (**Fig. 2*A*, *SI Appendix*, Fig. S2*D***).

### Functional categorization of screen results

Functional enrichment analysis (**Fig. 2*B*; Dataset S2**) revealed overrepresentation of pathways involved in DSB repair by homologous recombination and NHEJ, replication, chromosome dynamics, and protein degradation via vacuolar trafficking. We further validated screen hits by spot-testing deletion mutants on copper-supplemented plates (**Fig. 2*C*)**. We briefly summarize here the fully validated mutants. (See ***SI Appendix*, Discussion** for more detail, and ***SI Appendix*, Fig. S3** for mutants that had only mild Top2-hc sensitivity or failed final validation).

DNA repair mutants showed varied sensitivity to Top2-hc expression (**Fig. 2*C**; SI Appendix*, Fig. S3*A***). Homologous recombination mutants from the RAD52 epistasis group, including those encoding Mre11, Rad50, and Xrs2, displayed severe growth defects. NHEJ mutants showed more modest sensitivity, with *yku70* and *yku80* mutants exhibiting moderate defects while *dnl4*, *nej1*, *lif1*, and *pol4* mutants grew nearly normally. The STR complex mutants *top3* and *rmi1* conferred substantial sensitivity, while *sgs1* was even more sensitive, consistent with its additional Top3/Rmi1-independent functions in genome integrity (22). The *bud32* mutant, encoding a subunit of the EKC/KEOPS complex implicated in both tRNA modification and DNA repair (23), was the top scorer in validation tests.

DNA-protein crosslink (DPC) repair mutants demonstrated varied sensitivities (**Fig. 2*C*; *SI Appendix*, Fig. S3*A***). The *slx5* and *slx8* mutations, affecting a SUMO-targeted ubiquitin ligase complex, sensitized cells to Top2-hc, as did the absence of the SUMO E3 ligase Siz1, but not Nfi1. The Rtt101 E3 ubiquitin ligase complex mutants (*rtt101*, *mms1*, *mms22*, *ctf4*), recently implicated in removing topoisomerase-DNA covalent complexes (24), exhibited significant sensitivity. By contrast, neither *tdp1, wss1*, nor *ddi1* mutants showed obvious growth defects (***SI Appendix*, Discussion**).

Replication and chromosome segregation mutants affecting DNA polymerase δ (*pol32*), Okazaki fragment processing (*rad27*), replication fork protection (*csm3*, *mrc1*, *tof1*), and sister chromatid cohesion (*ctf18*, *ctf8*, *dcc1*) showed substantial sensitivity (**Fig. 2*C*; *SI Appendix*, Fig. S3*B***), suggesting that defects in repairing Top2-hc-generated damage are particularly lethal during S phase. Several centromere and kinetochore function mutants also displayed growth defects, potentially reflecting synergistic problems from combining replication-associated DNA damage with weakened chromosome segregation.

Additional sensitive mutants included vacuolar function mutants (e.g., *doa4*, *vps8*, and vacuolar H⁺-ATPase subunits); *fyv4* (mitochondrial ribosomal subunit); *ynr068c* (Rsp5 E3 ligase interactor); *brp1*; and *YLR358C/rsc2* (**Fig. 2*C*; *SI Appendix*, Fig. S3*E***).

### Comparison of Top1 and Top2 self-poisoning mutants

We compared screens using Top2-hc versus Top1-TA (18) (**Fig. 2*D***; **Dataset S3**; see ***SI Appendix,* Discussion** for more detail). The screens showed a significant but modest correlation (Spearman ρ = 0.339, p ≈ 7 × 10⁻¹²¹), indicating both shared and distinct pathway requirements. Of 385 genes with log₂ growth ratio ≥ 1 in either screen (control : mutant ratio), 23.9% were common hits (both screens), 29.4% were *TOP2-hc*-biased, and 46.8% were *TOP1-TA*-biased (**Fig. 2*D* and *E***). Of eleven TDA (<u>T</u>opoisomerase I <u>d</u>amage <u>a</u>ffected) genes previously identified in the Top1-TA screen, only *tda1* and *tda8* showed sensitivity to both Top2-hc expression and mAMSA treatment (***SI Appendix*, Fig. S3*H***).

Both screens identified core DNA repair pathways, particularly homologous recombination components (10 shared hits including *rad51, rad52, rad54, rad55, rad57, rad50, xrs2, top3, pol32, mus81*). Additional common hits included mutations affecting DSB repair, DNA damage response, sister chromatid cohesion, replication fork protection (*tof1, csm3, mrc1*), and kinetochore function (**Dataset S3**).

Top2-hc uniquely identified MAPK signaling components (8 hits) and the nucleolar Csm1/Lrs4 complex involved in rDNA segregation. Top1-TA specifically recovered DNA damage checkpoint machinery, including the Rad17-Mec3-Ddc1 clamp, its loader Rad24-RFC, checkpoint adaptor Rad9, sensor kinase Tel1, and endonuclease Sae2, plus Cul8 E3 ubiquitin ligase complex components Rtt101 and Rtt107 (**Dataset S3**).

Vesicle-trafficking and membrane-remodeling components were abundant in both screens. However, Top1-TA hits emphasized ESCRT-mediated multivesicular body (MVB) sorting, endosomal lipid signaling, and retromer/recycling pathways (21 specific hits), while Top2-hc hits instead highlighted pH control mechanisms (vacuolar ATPase subunits, Nhx1 exchanger) and vesicle tethering/fusion factors (CORVET/HOPS components, SNAREs) (**Dataset S3**).

### Human *TOP2A* and *TOP2B* hypercleavage mutations

Because the yeast R1128G single mutation alone yields hypercleavage behavior (19), we generated equivalent single human mutants (TOP2A-K1140G and TOP2B-K1158G, **Fig. 3*A**, SI Appendix*, Fig. S4*A***), designated TOP2A-hc and TOP2B-hc, and tested their toxicity in *top2-4* yeast (**Fig. 3*B***). Neither wild-type nor mutant TOP2A caused toxicity in *RAD52* and *rad52* strains at permissive temperature (**Fig. 3*Bi***). Both versions of TOP2A also supported *RAD52* growth at restrictive temperature (**Fig. 3*Bii***), confirming cross-species complementation consistent with prior studies (25, 26) and confirming that the TOP2A-hc mutant retains function. However, growth was slightly impaired with TOP2-hc expression compared to wild-type TOP2A, demonstrating a modest fitness cost of the hypercleavage mutant protein when it is the sole source of topoisomerase II activity in DSB repair-proficient yeast (**Fig. 3Bii**). Wild-type TOP2A also supported *rad52* growth but TOP2A-hc did not (**Fig. 3*Bi* and *ii***), consistent with the expected hypercleavage behavior.

**Fig. 3.**
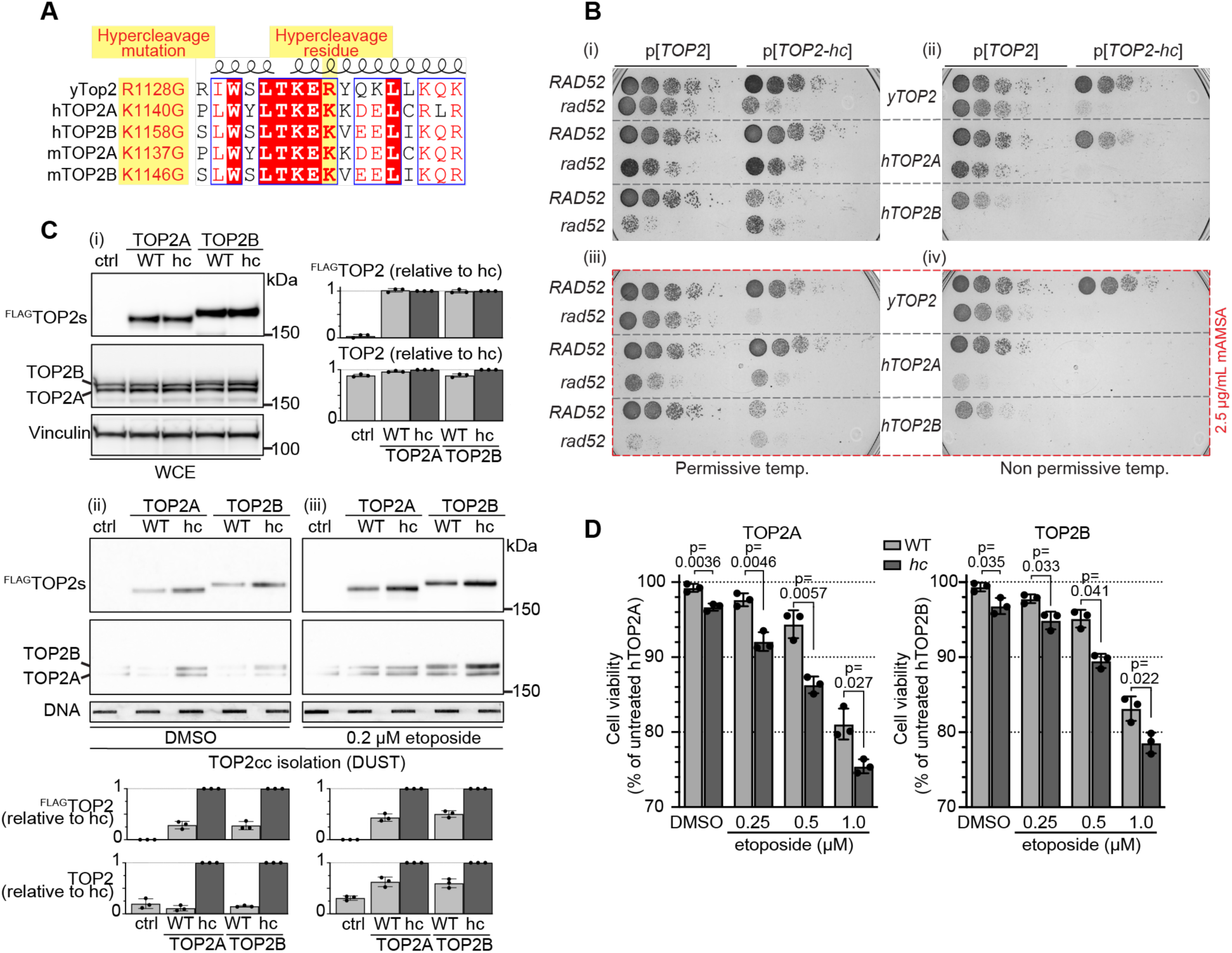
Generation and validation of human *TOP2-hc* alleles. (A) Alignment of a portion of the C-gate region of yeast Top2 with human and mouse TOP2A and TOP2B, highlighting the conserved lysine residues substituted by glycine in the hypercleavage mutants. (B) Complementation of temperature-sensitive *top2-4* by yeast and human *TOP2* alleles. Plasmids expressing WT or hypercleavage yeast Top2 or human TOP2A or TOP2B were transformed into *RAD52* or *rad52* strains in a *top2-4* background. Fivefold serial dilutions were spotted onto selective medium plates supplemented with 100 µM CuSO_4_ and incubated at permissive (28 °C, i and iii) or non-permissive (35 °C, ii and iv) temperatures, without (i and ii) or with (iii and iv) low-dose mAMSA. (C) Hypercleavage alleles increase covalent TOP2cc in RPE-1 cells. Immunoblots probed with anti-FLAG and anti-total-TOP2 antibodies are shown for whole-cell extracts (WCE, i) and DUST assays (ii, iii) after 24 h of DOX induction without (ii) or with (iii) additional co-treatment with 0.2 µM etoposide for 1 h. Anti-vinculin (i) and a slot-blot (2 µg DNA per slot) probed with anti-dsDNA antibody (ii, iii) serve as loading controls. Quantification of immunoblot chemiluminescence is expressed relative to internal controls (ctrl and TOP2A WT to TOP2A-hc, TOP2B WT to TOP2B-hc). (D) Induction of TOP2-hc confers cytotoxicity and enhances sensitivity to etoposide. Viability of RPE-1 cells expressing WT or hc TOP2A (left) or TOP2B (right) was tested following DOX induction and treatment with increasing concentrations of etoposide for 3 days. Viability is expressed as percentage of untreated WT controls (with DMSO, without DOX). Bars represent mean ± SD of three replicates; individual data points are shown. P values are from two-tailed Welch’s t-tests.

Low-dose mAMSA decreased *rad52* viability with wild-type TOP2A versus yeast Top2 (***Fig. 3Biii* and *iv***), suggesting either that human TOP2A is more mAMSA-sensitive (more covalent complexes generated) and/or that yeast Top2cc are repaired or tolerated better than human TOP2Acc. Viability in the presence of mAMSA decreased further with TOP2A-hc, and even *RAD52* failed to grow when TOP2A-hc was the sole topoisomerase II source at restrictive temperature (**Fig. 3*Biv***), confirming predicted hypersensitivity to poisons.

Wild-type TOP2B was tolerated in *RAD52* at permissive temperature and supported growth at restrictive temperature (25, 27), albeit less well than TOP2A (**Fig. 3*Bi* and *ii***). In contrast, TOP2B-hc caused diminished *RAD52* growth at permissive temperature and failed to complement *top2-4* at restrictive temperature (**Fig. 3*Bi* and *ii***), and it decreased *RAD52* growth in low-dose mAMSA at permissive temperature (**Fig. 3*Biii* and *iv***), consistent with both hypercleavage and hypersensitivity to poisons.

In *rad52*, wild-type TOP2B greatly diminished growth at permissive temperature and failed to complement at restrictive temperature (**Fig. 3*Bi* and *ii***), consistent with TOP2B forming DSBs readily even without poisons (28). Surprisingly, K1158G did not further decrease growth compared to wild-type TOP2B in *rad52* (**Fig. 3*Bi***), possibly due to selective pressure against plasmid-borne *TOP2B* expression.

### Toxicity of TOP2A-hc and TOP2B-hc in human cells

Triple-FLAG-tagged constructs introduced into human retinal pigment epithelial cells (RPE-1) yielded doxycycline (DOX)-inducible expression at levels comparable to endogenous (***SI Appendix*, Fig. S4*B***). DUST assays (detection of ubiquitylated-SUMOylated TOPcc (29)) showed a greater yield of TOP2cc for the hypercleavage versions in the absence of etoposide, confirming spontaneous hypercleavage activity, and TOP2cc yields increased further with a low dose of etoposide (**Fig. 3*C***). DOX induction of either TOP2A-hc or TOP2B-hc decreased viability and increased sensitivity to etoposide compared to uninduced cells, with TOP2A-hc conferring more etoposide sensitivity (**Fig. 3*D***). HeLa cells showed induced levels ∼2-3-fold compared to endogenous TOP2 (***SI Appendix*, Fig. S4*C***) accompanied by stronger viability effects: both TOP2A-hc and TOP2B-hc further decreased viability significantly in the absence of etoposide and substantially increased sensitivity to even low doses of etoposide **(*SI Appendix*, Fig. S4*D***).

### Hypercleavage mutations as screening tools in human cells

As proof of principle, we performed a short hairpin (sh)RNA dropout screen in HeLa using a library targeting 2028 genes enriched for involvement in DNA repair and DNA damage responses (**Dataset S4**). Cells with DOX-inducible expression of wild-type or hypercleavage TOP2 constructs below endogenous levels (***SI Appendix*, Fig. S4*E*)** were transduced with lentiviral shRNA library pools, then cultured for 5 passages over 20 days in the presence of DOX and a low dose (50 nM) of etoposide. This concentration was chosen to create selective pressure without overwhelming toxicity. shRNA abundance was quantified by deep sequencing, and z-scores were calculated to identify genes whose depletion enhanced etoposide sensitivity. At a threshold of z < -1.85, we identified 98 sensitizers for TOP2A-hc and 101 for TOP2B-hc, with 50 genes scoring strongly in both screens (**Fig. 4, Dataset S4**).

**Fig. 4.**
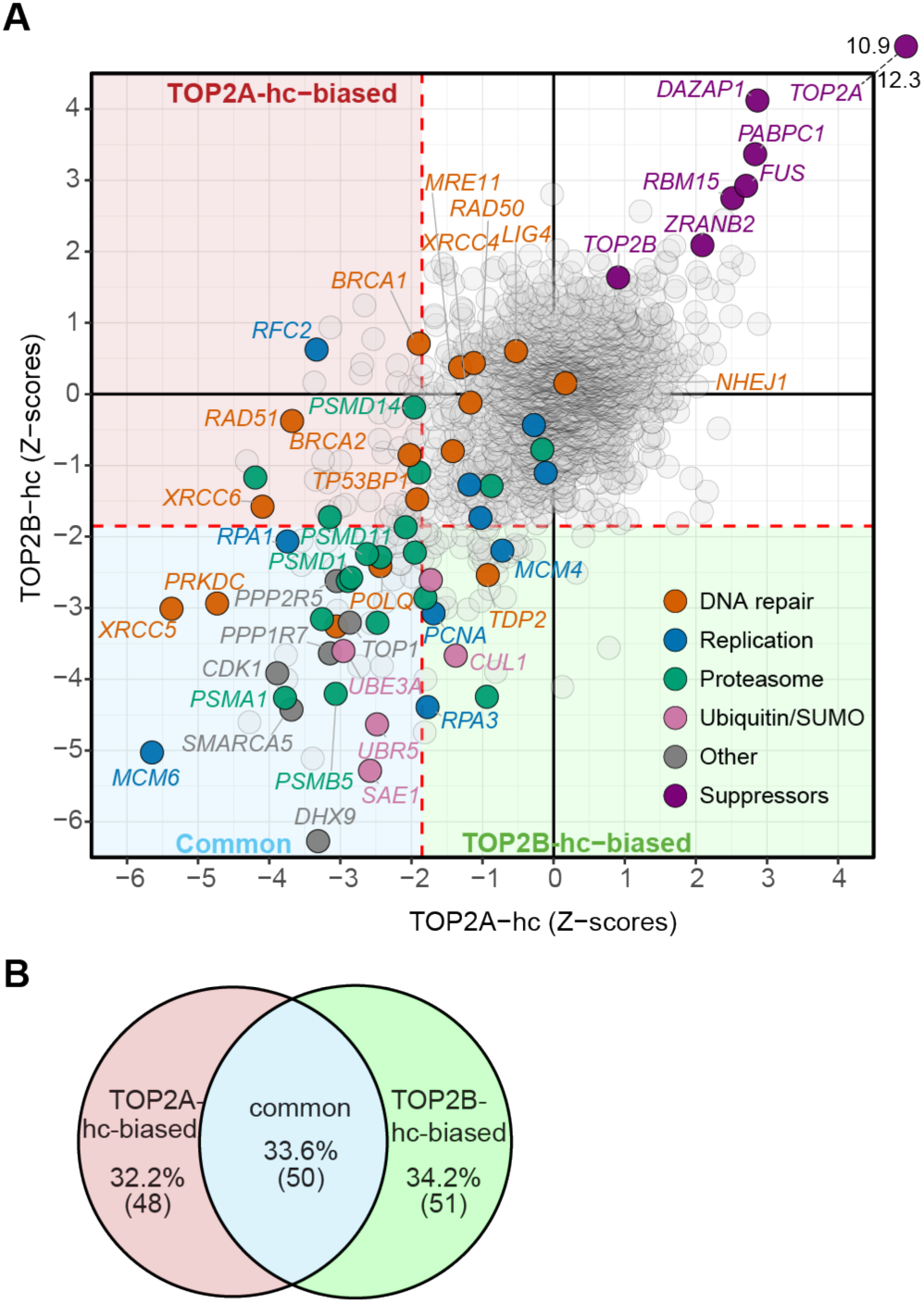
Comparative functional genomics of TOP2A-hc and TOP2B-hc reveals shared and isoform-specific vulnerabilities. (A) Comparative shRNA dropout results after expression of TOP2A-hc or TOP2B-hc. The scatterplot displays the control-anchored, pool-corrected z-scores, derived from log₂ fold-change values comparing the final (day 20, passage 5) to initial timepoints, for all genes in the screen. Each point represents the aggregated behavior of five independent shRNAs targeting a given gene. Negative z-scores indicate depletion (reduced viability), whereas positive z-scores indicate enrichment. Quadrant position and shading reflect differential impact on viability: the lower-left quadrant (z < –1.85 in both backgrounds; red dashed lines) denotes common sensitivity (blue), whereas preferential horizontal displacement (red, greater depletion with TOP2A-hc) or vertical displacement (green, greater depletion with TOP2B-hc) denotes isoform-biased effects. Selected genes are labeled, with functional categories color-coded as indicated. (B) Venn diagram summarizing overlapping and isoform-biased gene dependencies (color coded as for quadrant shading in panel A).

Unsurprisingly, several DSB repair factors required for recognition and processing of DNA ends—with or without covalent end blockage—were prominent hits (4, 30). For example, shRNAs targeting the DNA-PK complex components XRCC5 (Ku80), XRCC6 (Ku70), and PRKDC (DNA-PKcs) were strongly depleted in both screens, with a bias toward TOP2A-hc, with XRCC5 producing one of the most severe phenotypes in the screen (zA = -5.4, zB = -3.0), consistent with established roles in DSB end recognition, protection, and repair-pathway control (31). In contrast, downstream canonical NHEJ factors showed minimal or no effects (LIG4, XRCC4 and NHEJ1), suggesting that DNA-PK may contribute primarily to end protection or signaling rather than ligation *per se*. POLQ scored in both screens, implicating microhomology-mediated end joining (32) as an alternative to canonical end joining for repair of a subset of TOP2cc. Notably, the 5′-tyrosyl DNA phosphodiesterase TDP2 exhibited a pronounced TOP2B-hc-biased requirement, consistent with differential reliance on direct reversal of TOP2-DNA adducts between isoforms (4). In contrast, homologous recombination factors RAD51, BRCA2 and BRCA1 showed stronger requirements in the TOP2A-hc screen, consistent with their central roles in homologous recombination and replication-associated genome maintenance (33). Interestingly, shRNAs targeting MRE11 and RAD50 were not significantly depleted in either screen. This pattern may indicate that RAD51–BRCA-mediated strand invasion or replication-associated repair functions may be particularly important for viability under TOP2A-hc–induced stress, rather than full canonical MRN-dependent recombination.

Replication-associated factors were also hits. RPA subunits and the MCM2-7 helicase complex were required in both screens. MCM6 was the strongest single hit (zA = -5.6, zB = - 5.0), while other MCM subunits showed modest effects (MCM2, MCM3, MCM4, MCM7; MCM5 not in library). MCM6 may have unique functions beyond its role in the hexameric helicase, or it may simply be a more suitable shRNA target in this particular screen setup (e.g., allowing sufficient knockdown to give strong hypercleavage sensitivity without too much cell killing from the knockdown alone). PCNA showed a TOP2B-hc bias, but its loader RFC showed TOP2A-specific requirements, particularly RFC2. These findings are consistent with replication fork vulnerability in the presence of persistent TOP2cc (34). The RNA/DNA helicase DHX9 also scored as a sensitizer, potentially reflecting the interplay between transcription-associated R-loops and TOP2-mediated breaks (35).

Proteasome components were consistent, strong hits in both screens. Multiple 20S catalytic core subunits (PSMA and PSMB families) and 19S regulatory particle components (PSMD1, PSMD6 and PSMD11) were strongly depleted, supporting a central role for 26S proteasome-mediated processing of TOP2cc to permit exposure of DNA ends for downstream repair (6). Such proteolytic turnover is preceded by SUMO-and ubiquitin-dependent modification of TOP2cc, and components of these conjugation pathways likewise scored as sensitizers. The HECT-domain E3 ligases UBR5 and UBE3A and the SUMO E1 enzyme SAE1 were required in both isoform backgrounds, consistent with a SUMO–ubiquitin cascade that promotes proteasome-dependent degradation of TOP2cc (29), while the SCF scaffold CUL1 displayed a TOP2B-biased requirement, in agreement with CUL1-dependent ubiquitination of TOP2B (36).

Several additional strong sensitizers were shared, including the type I topoisomerase TOP1, the cell cycle regulator CDK1, multiple catalytic and regulatory subunits of the PP1 and PP2A phosphatase complexes, and the chromatin remodeler SMARCA5. The convergence of DNA topology factors, cell-cycle regulators, phosphatase signaling components, and chromatin remodeling machinery suggests that tolerance of TOP2cc requires coordinated control of DNA supercoiling, mitotic progression, and chromatin architecture (4, 37–39).

Suppressors of growth defects were also observed. TOP2A knockdown strongly enhanced survival in both screens (zA = +12.3, zB = +10.9), likely reflecting a correlation between TOP2A protein level and the formation of covalent complexes. The surprising strength of the effect in the TOP2B-hc screen, coupled with the minimal effect of TOP2B knockdown in either screen (zA = +0.9, zB = +1.6), indicates that endogenous TOP2A is a major driver of etoposide toxicity under these conditions. This isoform bias is consistent with prior work demonstrating that replication-associated conversion of TOP2cc into DSBs is predominantly TOP2A-dependent (12). Several RNA-processing factors, including DAZAP1, PABPC1, FUS, RBM15, and ZRANB2, also scored as suppressors. It is unclear whether these factors directly facilitate repair or tolerance of TOP2ccs, or if instead they work indirectly by regulating the expression of TOP2A or repair factors.

To contextualize our findings, we compared genetic dependencies in TOP2A-hc and TOP2B-hc screens with CRISPR sensitivity profiles from 31 DNA-damaging agents (8) (***SI Appendix* Fig. S4*F* and Dataset S5**). Spearman rank correlations were weak overall (|ρ| < 0.11), consistent with differences in screening platforms, cell lines, and genetic contexts. After Bonferroni correction (α = 0.0016), TOP2A-hc showed modest but significant positive correlations with several agents, including etoposide (ρ = 0.10, p = 8×10⁻⁶), cisplatin (ρ = 0.10, p = 4×10⁻⁵), and alkylating agents MNNG (ρ = 0.09, p = 2×10⁻⁴). In contrast, TOP2B-hc showed no significant correlations after multiple testing correction. These results suggest that TOP2A-mediated DNA damage is the more important contributor to sensitivity to genotoxic agents, in turn highlighting the potential of these hypercleavage constructs to dissect isoform specificity.

### Mouse models expressing *Top2A* and *Top2B* hypercleavage mutations

To establish TOP2-hc as an *in vivo* tool, we generated DOX-inducible transgenic mice expressing 2×FLAG-tagged mouse TOP2A-K1137G or TOP2B-K1146G (equivalent to human TOP2A-K1140G and TOP2B-K1158G (**Fig. 3*A***)). The system employs a dual-cassette design (40): *CAG-LSL-rtTA3* drives DOX-responsive transactivation, while *TRE-LSL-Top2-hc-IRES-EGFP* encodes the hypercleavage mutant (***SI Appendix* Fig. S5*A***). Cre-mediated excision of both *loxP*-stop-*loxP* (LSL) cassettes renders *Top2-hc* expression DOX-inducible. Using the germline deleter strain *E2a-Cre*, we achieved whole-body recombination and confirmed robust TOP2-hc-FLAG expression in multiple tissues, with the highest levels in liver, pancreas, and heart and undetectable levels in brain and testis (**Fig. 5*A*, *SI Appendix* Fig. S5*B***).

**Fig. 5.**
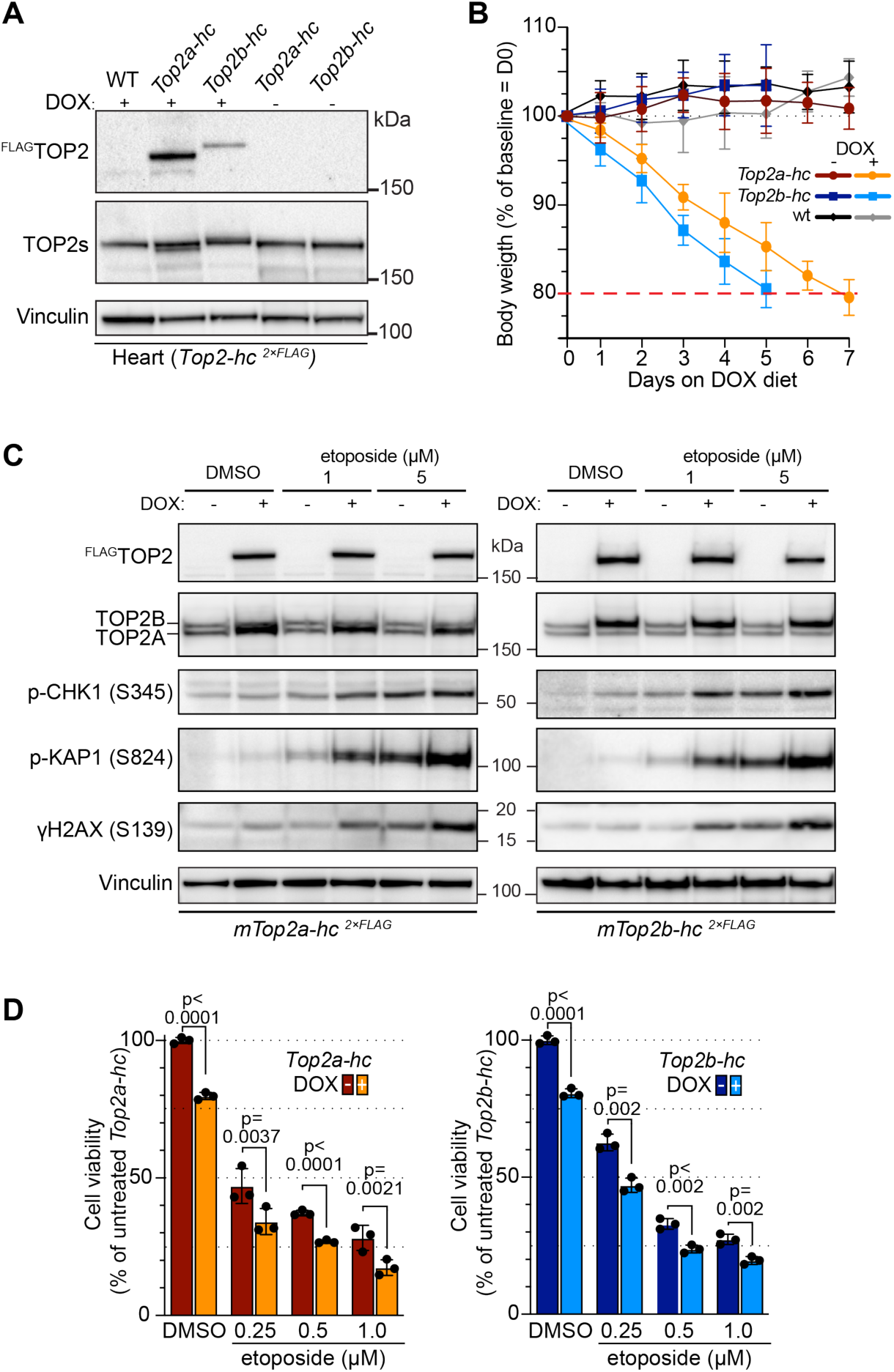
Generation and validation of murine *Top2-hc* alleles and mouse models. (A) DOX-inducible expression of FLAG-tagged TOP2A-hc or TOP2B-hc *in vivo*. An immunoblot of heart extracts is shown, probed with anti-FLAG or pan-TOP2 antibodies. Vinculin serves as a loading control. (B) Intrinsic toxicity of TOP2-hc expression *in vivo*. Body weights (percentage of baseline at day 0) are shown after start of a DOX-containing diet (625 mg/kg) or maintenance on regular chow. Body weight was recorded daily for 5 days (*Top2b-hc*) or 7 days (*Top2a-hc*), at which point most DOX-treated animals reached the humane endpoint of ∼20% weight loss (dashed red line). Error bars show mean ± SD (n = 5 mice per genotype). (C) DNA damage response in immortalized MEFs expressing TOP2-hc. Immunoblots are shown of whole cell extracts prepared from with or without DOX-induced expression of TOP2A-hc (left) or TOP2B-hc (right), co-treated with DMSO (control) or etoposide (1 or 5 µM for 90 min). Additional blots and quantification are shown in ***SI Appendix*, Fig. S6*A***. (D) Intrinsic cytotoxicity and enhanced sensitivity to etoposide from TOP2-hc expression in immortalized MEFs. Cell viability was assessed as in Fig. 3D for MEFs isolated from transgenic mice expressing TOP2A-hc (left) or TOP2B-hc (right) following DOX induction and treatment with increasing concentrations of etoposide for 4 days. Viability is expressed as percentage of untreated, uninduced controls (with DMSO, without DOX). Bars represent mean ± SD of three replicates; individual data points are shown. P values are from two-tailed Welch’s t-tests.

DOX treatment of young adult mice caused progressive weight loss over 7 days, with *Top2b-hc* proving more toxic than *Top2a-hc* (**Fig. 5*B***), demonstrating that hypercleavage mutations are deleterious *in vivo*. DOX-treated TOP2A-hc animals revealed pallor of the liver, darker intestines and reduced spleen size relative to non-induced littermates (***SI Appendix* Fig. S5*C*)**. The involvement of intestine, a highly proliferative tissue, supports a replication-dependent mechanism of toxicity, but the prominent liver and spleen phenotype indicates that toxicity is not restricted to actively cycling tissues. Similar gross changes were observed following induction of TOP2B-hc (***SI Appendix* Fig. S5*C***), indicating that tissue-selective injury reflects persistent TOP2cc formation *per se* rather than isoform-specific effects.

Immortalized embryonic fibroblasts derived from these mice showed strong, specific DOX induction of FLAG-TOP2-hc, resulting in an approximately twofold increase in total TOP2 signal (**Fig. 5*C* and SI Appendix Fig. S6*A***). The FLAG-TOP2-hc was localized to the nucleus as expected and resulted in elevated γH2AX (**SI Appendix Fig. S6*B***), as well as other markers of DNA damage—phosphorylation of ATM, CHK1, 53BP1, and KAP1 (**Fig. 5*C* and SI Appendix Fig. S6*A***). Etoposide treatment further enhanced these DNA damage responses (**Fig. 5*C* and SI Appendix Fig. S6*A***). DUST assays demonstrated TOP2cc formation in the absence of etoposide, confirming the intrinsic hypercleavage behavior, and etoposide treatment increased TOP2cc recovery in a dose-dependent manner (**SI Appendix Fig. S6*C***). In DOX-induced samples, the DUST-recovered signal was dominated by the TOP2-hc-FLAG isoform, while non-induced controls at the same etoposide doses recovered both endogenous TOP2 isoforms without bias. DOX induction strongly shifted the recovered complexes toward the mutant species, indicating that TOP2-hc is overrepresented among trapped complexes and consistent with the idea of preferential trapping of the mutant enzyme (**SI Appendix Fig. S6*C***). Moreover, cell viability assays demonstrated DOX-induced toxicity that was further enhanced by etoposide in a dose-dependent manner (**Fig. 5*D***).

## Discussion

Genetic dissection of cellular responses to topoisomerase-mediated DNA damage has been limited by reliance on pharmacological poisons, which cause off-target effects and— critically for mammalian systems—target both TOP2A and TOP2B (41, 42). Here we establish an inducible, cross-species toolkit of TOP2 hypercleavage mutants, providing the first genetically defined systems for isoform-specific investigation of TOP2cc biology.

The proof-of-principle screens in yeast and human cells served primarily to validate that TOP2-hc mutants generate expected TOP2cc-engaging repair pathways. Both screens recovered core homologous recombination factors, proteasome components, and DNA-PK—all established players in TOP2cc processing (4, 6)—confirming that hypercleavage mutations produce lesions requiring canonical repair machinery (19). The screens also identified unexpected hits (vesicle trafficking in yeast, RNA-processing factors in human cells), but these should be considered hypothesis-generating rather than definitive at present. In particular, the shRNA approach, with its targeted gene list, lacked functional validation.

The value of these screens lies in demonstrating that TOP2-hc mutants are suitable for large-scale genetic interrogation: they establish appropriate expression levels, confirm that growth/viability defects are within screening dynamic range, and show that combining TOP2-hc with low-dose poison amplifies selective pressure. Future genome-wide CRISPR screens in multiple cell types, with rigorous validation and mechanistic follow-up, should define the full spectrum of repair requirements and resolve isoform-specific dependencies.

The mammalian TOP2 paralogs have proven difficult to study independently: they share ∼70% sequence identity, localize to overlapping compartments, and are targeted by the same poisons (5, 43). Knockout or knockdown approaches are confounded by compensation and pleiotropic effects (5), while poison-based chemical genetics cannot distinguish isoform contributions. TOP2A-hc and TOP2B-hc provide a powerful genetic solution, allowing controlled generation of isoform-specific TOP2cc without perturbing endogenous TOP2 levels. This separation of cleavage-complex formation from protein abundance provides a conceptual and experimental solution to a long-standing barrier in the field.

Although exploratory, the shRNA screen provided proof-of-principle that differential genetic dependencies can be detected. This capability enables questions previously intractable with existing tools: Which repair pathways show absolute isoform preference? What determines tissue-selective toxicity of TOP2 poisons? We note, however, that the screen was performed in the presence of low-dose etoposide, which would be expected to also poison the endogenous TOP2A and TOP2B, albeit to a lesser extent than the transgenically expressed TOP2-hc. Thus, the resulting hits should be interpreted as isoform-biased genetic interactions under combined TOP2-hc and endogenous TOP2 poison stress, rather than as strictly isoform-specific dependencies.

We also note that there are multiple possible reasons why the TOP2A-hc and TOP2B-hc mutants might differ from one another in their biological effects, including isoform-specific differences in the degree to which the mutations promote DSB formation, differences in the genomic and cell cycle context of DSB formation, and differences in repair efficiency. Biochemical analysis of purified human or murine TOP2A-hc and TOP2B-hc proteins may be helpful in discriminating some of these possibilities.

In keeping with prior work (19, 44), TOP2-hc mutants confer marked hypersensitivity to otherwise well-tolerated poison doses across all systems tested. This property positions them as sensitized backgrounds for identifying new topoisomerase inhibitors and rapidly profiling isoform selectivity—capabilities particularly relevant given interest in developing TOP2A-selective or TOP2B-sparing agents to reduce cardiotoxicity (45). The hypersensitivity also validates the underlying mechanism: hypercleavage mutations shift the cleavage-religation equilibrium as intended, and poisons act synergistically by further stabilizing the cleavage complex.

The DOX-inducible mouse models demonstrate that *TOP2-hc* alleles can be induced in adult tissues, generate measurable DNA damage, and sensitize cells to etoposide. However, because these alleles are expressed from an inducible transgenic system rather than the endogenous loci, their tissue, cell-type, and cell-cycle distributions do not reflect the physiological expression patterns of the two isoforms. Thus, these models are best interpreted as tools to induce defined TOP2cc burdens *in vivo*, rather than as faithful models of endogenous isoform-specific TOP2 biology. Nonetheless, these models establish feasibility for future studies using tissue-specific Cre drivers to model tissue-specific vulnerabilities to TOP2cc during development, and investigate questions inaccessible to cell culture, e.g., whether cardiac-specific TOP2B-hc can reproduce doxorubicin cardiotoxicity (45).

In summary, the TOP2 hypercleavage toolkit establishes a genetically defined strategy for generating and studying isoform-specific TOP2cc across model systems. By enabling inducible stabilization of cleavage complexes in yeast, cultured mammalian cells, and mice, this work provides a foundation for mechanistic dissection of TOP2cc formation, processing, and biological outcome in contexts that were previously inaccessible. The principal impact of this study is not the specific pathways identified here, but the experimental framework it creates for resolving isoform-specific biology and tissue-selective vulnerability in topoisomerase-mediated DNA damage.

## Materials and Methods

### Plasmids

Plasmids generated in this study are listed in **Dataset S6**, including Addgene plasmid IDs for deposited yeast, mouse, and human wild-type and hypercleavage TOP2 expression constructs. Details of plasmid construction are provided in ***SI Appendix, Materials and Methods***.

### Yeast methods

*Saccharomyces cerevisiae* strains are listed in **Dataset S7**. Strains for the complementation assays carrying the temperature-sensitive *top2-4* were JN362a t2-4 and JN394a t2-4 *rad52::LEU2* (21). All other strains were in the BY4741 background (46). Gene deletions and C-terminal tagging of *RAD52* with GFP were made using a PCR-based approach (47, 48).

Methods for spot testing yeast growth are provided in ***SI Appendix, Materials and Methods***. The SPA screen was performed essentially as described (18), with plasmids expressing *TOP2* or *TOP2-hc*, using the *MATa* yeast gene disruption library from Open Biosystems (49). Screening was performed under basal and copper-induced expression conditions. Details are provided in ***SI Appendix, Materials and Methods***. For functional enrichment analysis, data were obtained from the *Saccharomyces* Genome Database (50) and analyzed using ShinyGO v0.82-0.85 (51).

For immunoblotting, trichloroacetic acid (TCA) yeast cell extracts were prepared from 5– 10 ml of exponentially growing cultures in SC-LEU. Fluorescence microscopy was performed to visualize GFP-tagged proteins (Top2 and Rad52). Top2cc isolation was performed using the ICE assay, as described (52). Details for immunoblotting, microscopy, and ICE assays are provided in ***SI Appendix, Materials and Methods***. Primary and secondary antibodies are listed in **Dataset S8.**

### Mammalian methods

Human *TOP2* expression constructs were introduced into HeLa Flp-In™ T-REx™ cells (Thermo Fisher Scientific, R71407) by targeted recombinase insertion, and into RPE-1 cells through lentiviral transduction. MEFs were isolated from E13.5 mouse embryos and immortalized by lentiviral transduction with SV40 large T antigen. DUST assays were performed as described (29) with modifications. Cell viability following DOX-induction of TOP2 expression and etoposide treatment was determined using CellTiter-Blue reagent (Promega) and a SpectraMax iD5 microplate reader (Molecular Devices). Viability was normalized to untreated wild-type controls within each experiment. Cell lysates were prepared in SDS sample buffer, sonicated briefly, boiled, clarified by centrifugation, and analyzed by SDS-PAGE on NuPAGE Bis-Tris or Tris-acetate gels followed by transfer to PVDF membranes, blocked, probed with primary and secondary antibodies, and developed by chemiluminescence. Immunofluorescence was performed on MEFs using standard fixation, permeabilization, and antibody staining procedures, followed by DAPI counterstaining and imaging on a Zeiss Axio Observer Z1 Marianas Workstation. Details of the derivation and culture of cell lines, the DUST and cell viability assays, immunoblotting, and immunofluorescence are provided in the ***SI Appendix, Materials and Methods***.

For the shRNA screen, the HeLa Flp-In™ T-REx™ cell lines were transduced with a pooled shRNA library targeting the human DNA damage response, prepared by the MSK Genome Editing and Screening Core Facility. This library contained 10,607 shRNA constructs across 6 subpools, targeting 2,028 unique human protein-coding genes with 5 shRNA constructs per gene, along with negative controls targeting firefly and Renilla luciferases, cloned in SGEP-based miR-E lentiviral vectors (53). Transductions were performed at low multiplicity of infection (∼0.2) followed by puromycin selection. Baseline samples (T0) were collected after selection, and cells were cultured for 20 days (5 passages) under DOX induction (2 µg/mL) and low-dose etoposide (50 nM). Integrated shRNA cassettes were recovered from genomic DNA by PCR and quantified by Illumina sequencing. Read counts were normalized and summarized to gene-level log₂ fold-changes relative to T0. Gene scores were converted to z-scores within each subpool and condition, followed by normalization using shared internal controls and correction for subpool-specific offsets while preserving condition effects. Hits were defined using a z ≤ −1.85 threshold corresponding to an estimated false-positive rate of ≤1%. Analyses were performed in R (v4.3.1, R core team 2023). Detailed library composition, sequencing procedures, and statistical pipeline are described in ***SI Appendix, Materials and Methods* and Dataset S4.**

### Mice

Experiments conformed to the US Office of Laboratory Animal Welfare regulatory standards and were approved by the MSK Institutional Animal Care and Use Committee (protocol number 01-03-007). Mice were euthanized by CO_2_ asphyxiation prior to tissue harvest. Cre-conditional, DOX-inducible *Top2-hc* knock-in mouse models were generated using FlpE-mediated recombinase-mediated cassette exchange (RMCE) in mouse embryonic stem cells (mESCs). cDNAs encoding *mTop2a-hc-2×FLAG* or *mTop2b-hc-2×FLAG* were cloned into the *Col1a1* targeting vector (cTGM) and electroporated into KH2 mESCs to achieve single-copy transgene insertion downstream of the endogenous *Col1a1* locus (54, 55). Genotyping was performed by Transnetyx Inc. (Cordova, TN, USA) using real-time PCR. Detailed methods for animal husbandry, generation of knock-in lines, and extraction of protein for immunoblotting are provided in ***SI Appendix, Materials and Methods***.

### Generative AI use

ChatGPT (version 5.2, OpenAI) was used to assist in exploratory analysis of shRNA screen data and in drafting R scripts for data visualization. Claude 4 Sonnet (Anthropic, San Francisco, CA, USA) was used to edit portions of the manuscript for clarity and grammar. After using these tools, the authors reviewed and edited the content for accuracy and take full responsibility for the content of the published article.

## Author contributions

D.O., M.M., and S.K. conceived the research. M.M. constructed plasmids and carried out the SPA screen with J.D. under supervision from R.J.D.R and R.R. J.L.N. provided unpublished data and the yeast *TOP2-hc* mutant prior to publication. D.O. performed all other experiments. D.O. and S.K. wrote the manuscript.

## Acknowledgments

This article is subject to HHMI’s Immediate Access to Research policy, which requires that this article be made publicly available as initial and revised preprints deposited on a designated preprint server under a CC BY 4.0 license.

We are grateful to James Wang (Harvard University) for providing polyclonal antibodies against yeast Top2; members of Rothstein laboratory for help with the SPA screen; Xiaolan Zhao (MSK) for yeast strains from the knockout collection; Mathew Jones (Jallepalli laboratory, MSK) for assistance with the shRNA screen, providing cell lines and cell culture advice; Ralph Garippa, Sanjay Mehta, and Daniel Zakheim (MSK RNAi & Gene Editing Core Facility) for providing the shRNA library and for assistance with the shRNA screen; Ambereen Khan and Zhen Zhao for transgene targeting in mESCs and Direna Alonso-Curbelo for providing plasmids and advice on mouse experiments (Scott Lowe laboratory, MSK); Sang Yong Kim for the mouse embryonic stem cells injections into blastocysts (NYU Langone Health); the MSK Molecular Cytogenetics Core Facility for karyotyping; Pedro San-Segundo (IBFG, Salamanca) for sharing plasmids; and members of the Keeney laboratory, especially Mariko Sasaki for plasmid construction and Soonjoung Kim for discussions. MSK core facilities are supported by National Cancer Institute cancer center support grant P30 CA08748. This work was supported by NIH grants R01 GM058673 (to S.K.), R35 GM118092 (to S.K.), R03 CA297372 (to J.L.N.), and R35 GM118180 (to R.R); Grant I12-0008 from the Starr Cancer Consortium (to Jessica Tyler and S.K.), the former Center for Metastasis Research (since incorporated into the Alan and Sandra Gerry Metastasis & Tumor Ecosystems Center; to S.K.).

