## Supplementary material for "A multi-species toolkit of *TOP2* hypercleavage mutants for studying topoisomerase II–mediated DNA damage": SI Appendix

This pdf contains:
Supplemental Discussion
Supplemental Methods
Supplemental References
Supplemental Figures S1-S6

Additional information, in separate .xlsx files:

**Dataset S1**. Full, validation and hits yeast *TOP2-hc* screen results
**Dataset S2**. Functional enrichment yeast *TOP2-hc* hits.
**Dataset S3**. Yeast SPA screens comparison: TOP2hc vs TOP1TA.
**Dataset S4**. shRNA results.
**Dataset S5**. shRNA vs CRISPR.
**Dataset S6**. List of plasmids.
**Dataset S7**. List of yeast strains.
**Dataset S8**. List of antibodies.

### Supplemental Discussion

#### Functional categorization of yeast Top2-hc screen results – Supplemental Details

**DNA repair pathways.** Building on earlier observations (19) (**Fig. 1*E***), numerous DSB repair-deficient mutants exhibited impaired growth when expressing Top2-hc (**Fig. 2*C*; *SI Appendix*, Fig. S3*A***). The most pronounced vulnerabilities occurred in mutants from the *RAD52* epistasis group, which function in homologous recombination pathways (56). This group includes genes encoding the Mre11-Rad50-Xrs2 (MRX) complex, which initiates DSB resection and participates in multiple repair processes.

By comparison, mutants defective in NHEJ exhibited more variable and generally milder phenotypes. Deletion of genes encoding the Ku heterodimer (*yku70*, *yku80*), which binds DSB ends and recruits downstream repair factors, produced moderate growth defects that were nonetheless less severe than those observed with homologous recombination mutants. Other NHEJ pathway components—*dnl4* (DNA ligase IV), *nej1* (NHEJ accessory factor), *lif1* (Dnl4 cofactor), and *pol4* (DNA polymerase involved in end processing)—showed essentially wild-type growth levels when challenged with Top2-hc expression (**Fig. 2*C***; ***SI Appendix*, Fig. S3*A***).

The STR complex—comprising Sgs1 (RecQ helicase), Top3 (topoisomerase III), and Rmi1—facilitates repair of replication-associated DSBs and other forms of DNA damage (57). Sgs1 also has additional functions independent of Top3 and Rmi1, particularly in DSB end resection (22, 58). Mutants lacking Top3 or Rmi1 displayed substantial Top2-hc sensitivity, while the *sgs1* deletion conferred even greater sensitivity (**Fig. 2*C***), consistent with its broader role in maintaining genome stability.

Spot tests further validated the remarkable Top2-hc sensitivity of *bud32*, which emerged as the highest-scoring hit in our validation screen (**Fig. 2*C***; **Dataset S1**). Bud32 functions as both a protein kinase and ATPase within the EKC/KEOPS complex, primarily known for its essential role in tRNA modification but recently implicated in homologous recombination-mediated DNA repair (23).

**Repair pathways for covalent DNA-protein crosslinks.** Post-translational modifications by SUMOylation and ubiquitination followed by proteolytic degradation via the 26S proteasome facilitate removal of trapped Top2cc (59, 60). The Slx5-Slx8 SUMO-targeted ubiquitin ligase (STUbL) complex recognizes previously SUMOylated Top2cc and marks it for proteasomal turnover (29, 61). Consistent with this mechanism, both *slx5* and *slx8* deletions sensitized cells to Top2-hc expression (**Fig. 2*C***). Loss of Siz1, a SUMO E3 ligase whose activity precedes STUbL-mediated degradation (29), produced comparable sensitivity (**Fig. 2*C***). However, deletion of Nfi1 (also called Siz2), another SUMO E3 ligase with overlapping functions, did not confer Top2-hc sensitivity (**Fig. 2*C***).

In *S. cerevisiae*, tyrosyl-DNA phosphodiesterase 1 (Tdp1) catalyzes hydrolysis of both 3′- and 5′-phosphotyrosyl linkages in Top1cc and Top2cc, as yeast lacks a TDP2 ortholog specific for 5′ bonds (62, 63). While *tdp1* showed synthetic sickness in initial full-screen plates (***SI Appendix*, Fig. S2*C***), subsequent spot test validation revealed no clear growth defect with Top2-hc expression relative to control cells overexpressing wild-type Top2 (***SI Appendix*, Fig. S3*A***). We additionally examined Wss1, the yeast ortholog of human SPRTN, which is a SUMO-targeted metalloprotease that removes SUMOylated chromatin-associated proteins (64, 65), and Ddi1, a protease contributing to Top1cc repair in yeast (66). Neither *wss1* nor *ddi1* were identified in the screen, and both failed to show growth defects in spot test validation (***SI Appendix*, Fig. S3*A***).

Rtt101 (alternatively designated Cul8) is an E3 ubiquitin ligase that forms a complex with Mms1, Mms22, and Ctf4 (67) to promote replication-associated DNA repair and facilitate replication fork progression through DNA lesions (68). Recent work has implicated Rtt101 and its interacting partners in the removal of Top1cc, Top2cc, and other DPCs, with corresponding deletion mutants displaying sensitivity to low-dose camptothecin or etoposide treatment (24). Our results align with these findings, as *mms22* and *ctf4* mutants were picked up in the initial and validation screens and *rtt101* and *mms1* also showed marked sensitivity to Top2-hc expression when tested directly in spot tests (**Fig. 2*C***).

**DNA replication, sister chromatid cohesion, and kinetochore assembly.** Multiple mutants compromised in replication stress responses or replication-coupled DNA damage repair displayed significant Top2-hc sensitivity. These included deletions affecting Pol32, a non-essential subunit of DNA polymerase δ involved in break-induced replication and other repair processes (*pol32*) (69); Rad27, a 5'-flap endonuclease critical for Okazaki fragment processing (*rad27*) (70); and components of the replication fork protection complex that prevent fork collapse during replication stress (*csm3*, *mrc1*, *tof1*) (71) (**Fig. 2*C***).

Additional sensitive mutants included *rtt109* and *asf1*, encoding a histone acetyltransferase and nucleosome assembly factor, respectively, both required for chromatin reassembly and recovery following DSB repair (18, 72, 73). Additional chromatin assembly/regulation mutants (*swi4, swi6, snf5, swi3, yaf9, spt2, spt21, sin3, hda3, swd1*) also scored in the validation screen (***SI Appendix*, Fig. S3*D***), although these were not subsequently tested by spot growth assays. We also identified subunits of an RFC-like alternative clamp loader (*ctf18*, *ctf8*, *dcc1*) that functions in establishing sister chromatid cohesion during DNA replication (74) (**Fig. 2*C***). The collective sensitivity of these replication-related mutants suggests that failure to properly repair Top2-hc-induced DNA damage becomes especially detrimental during S phase.

Synthetic growth defects also emerged with mutations disrupting various aspects of centromere structure and kinetochore assembly. Affected mutants included *cbf1*, encoding a sequence-specific centromere DNA-binding protein (**Fig. 2*C***); components of the Ctf19 complex involved in kinetochore assembly (*ctf19*, *mcm21*, *chl4*); and outer kinetochore components (*ctf3*, *mcm16*, *mcm22*) (75-77) (***SI Appendix*, Fig. S3*B***). The mechanistic basis for these sensitivities remains to be fully elucidated, but we hypothesize that combining replication-associated DNA damage from Top2cc with compromised mitotic chromosome segregation creates synergistic defects that impair cell viability.

**Vacuolar and vesicle trafficking pathways.** Several mutations affecting vacuolar function and intracellular vesicle trafficking conferred varying degrees of growth impairment upon Top2-hc expression (**Fig. 2*C***; **SI Appendix, Fig. S3*E***). Examples include *doa4*, encoding a deubiquitinating enzyme involved in protein sorting, and *vps8*, encoding a component of the CORVET complex required for vacuolar protein sorting (78). Additionally, deletions of genes encoding subunits of the vacuolar $H^+$-ATPase complex (*vma1*, *vma8*, *vma16*) caused Top2-hc sensitivity. These mutations disrupt vacuolar acidification, leading to defects in protein degradation, autophagy, and various stress response pathways (79-81).

**Additional genes with diverse functions.** Beyond the major functional categories, several other deletions produced notable Top2-hc sensitivity. *FYV4* deletion rendered cells extremely vulnerable to Top2-hc expression (**Fig. 2*C***). Fyv4 is a component of the mitochondrial small ribosomal subunit, and its loss impairs respiratory growth while increasing susceptibility to environmental stresses (82, 83).

*YNR068C* (*BUL3*) deletion also had a pronounced effect (**Fig. 2*C***). Ynr068c interacts with Rsp5, a NEDD4 family E3 ubiquitin ligase, and this interaction is required for cellular survival following treatment with the DNA-damaging agent phleomycin; the Rsp5 pathway likely functions in ubiquitin-mediated sorting of plasma membrane proteins (84).

The *brp1* mutant exhibited enhanced Top2-hc sensitivity as well (**Fig. 2*C***). Brp1 plays a role in degrading the membrane protein Agp2 upon cycloheximide treatment, though the precise mechanism remains incompletely understood (85).

Finally, deletion of *YLR358C*, a verified gene of unknown function, caused substantial Top2-hc sensitivity (**Fig. 2*C***). This gene partially overlaps in antisense orientation with *RSC2*, which encodes a well-characterized component of the RSC chromatin remodeling complex (86). Since *rsc2* deletion produces similar Top2-hc sensitivity (**Fig. 2*C***), it remains unclear whether one or both genes contribute to Top2-hc tolerance.

#### Comparison of yeast screens with Top1 and Top2 self-poisoning mutants – Supplemental Details

To understand the similarities and differences in cellular responses to distinct topoisomerase lesions, we compared gene products involved in counteracting self-poisoning by Top2-hc versus Top1-TA mutants (18) (**Fig. 2*D***, **Dataset S3**). Statistical analysis revealed a significant correlation between the two SPA screen profiles (Spearman $\rho = 0.339$, $p \approx 7 \times 10^{-121}$), suggesting partial overlap between the pathways recruited by each type of trapped topoisomerase complex. This indicates that while certain core cellular mechanisms respond to both lesion types, each also requires specialized factors. We further identified 385 unique genes (8.77% of 4,389 tested) that showed $\log_2$ growth ratio $\leq -1$ (topo mutant : EV) in one or both screens (positioned below and/or to the left of the dashed red lines in **Fig. *2D*** and detailed in **Dataset S3**).

**Shared requirements.** Both screens highlighted the fundamental importance of DNA repair pathways, with particular dependence on homologous recombination. Ten mutants identified as shared hits in this category included core recombination components: *rad51* (recombinase), *rad52* (mediator), *rad54* (chromatin remodeler), *rad55* and *rad57* (Rad51 paralogs), *rad50* and *xrs2*, *top3*, *pol32*, and *mus81* (structure-selective endonuclease).

Beyond the core HR machinery, common hits included numerous factors categorized under gene ontology terms for DSB repair (21 genes) and broader DNA damage response (31 genes), such as *rad27* (flap endonuclease), *rtt109* (histone acetyltransferase), and *asf1* (histone chaperone). Genes involved in sister chromatid cohesion (n=15) and replication fork management (n=8) were also shared between screens. Kinetochore function genes (n=9) including *ctf19, mcm16, mcm21, chl4, chl1, mcm22, ctf3,* and *cbf1* (87, 88) were identified in both screens. Fork protection complex mutants *tof1, csm3*, and *mrc1*, along with mutations affecting the Ctf18-RFC complex (*ctf18*, *ctf8*, and *dcc1*), were also common to both screens, as were members of the Cul8 E3 ubiquitin ligase complex (*ctf4* and *mms1*), plus the spot-test validated *rtt101* and *mms1*. These convergent findings demonstrate that both types of topoisomerase-mediated lesions ultimately depend on recombination-mediated repair mechanisms and proper regulation of replication fork dynamics and sister chromatid cohesion.

**Top2-hc-specific requirements.** The Top2-hc screen uniquely identified eight mutations affecting mitogen-activated protein kinase (MAPK) signaling pathways, including *swi4, cln1, sic1, bmh1*, and *ste11*. MAPK pathways enable eukaryotic cells to mount responses to diverse environmental signals, ranging from hyperosmotic stress to chemical challenges such as arsenite exposure or treatment with the DNA alkylating agent methyl methanesulfonate (89-91)

Additionally, Top2-hc specifically identified the nucleolar Csm1/Lrs4 complex (**Dataset S3**), which contributes to rDNA silencing, recruitment of condensin to replication fork barrier sites within the rDNA array, and proper segregation of rDNA repeats during cell division (92, 93).

**Top1-TA-specific requirements.** The Top1-TA screen uniquely recovered mutations affecting critical components of the DNA damage checkpoint machinery. These included the PCNA-like clamp complex Rad17-Mec3-Ddc1 and its loader Rad24-RFC, which together load the clamp onto 5′-recessed DNA junctions to initiate checkpoint signaling cascades (94, 95). The checkpoint adaptor protein Rad9, which undergoes Mec1/Tel1-dependent phosphorylation to couple the clamp signal to Rad53 activation (96), was also Top1-TA-specific. Additional specific hits included the MRX-accessory factor Sae2, which promotes resection initiation, and sensor kinase Tel1, which is recruited to unresected DNA ends via the MRX complex (97).

**Vesicle trafficking and membrane remodeling.** Numerous vesicle-trafficking and membrane-remodeling components were required for cellular responses to topoisomerase-mediated stress, consistent with their established roles in maintaining genome integrity and activating responses to genotoxic stress (98-100). Endosomal proteins were the most abundant category, with 21 Top1-TA-specific hits, 8 Top2-hc-specific hits, and 10 common hits.

The Top1-TA-specific hits showed enrichment for ESCRT-mediated multivesicular body (MVB) sorting and endosomal lipid signaling pathways. These included ESCRT-I/II/III factors and adaptors (*stp22, mvb12, vps28, snf8, vps20, vps60, did2, srn2*), the Vps30-Vps38 PI3K complex, retromer and recycling components (*vps29, rcy1*), and coat/adaptor proteins (*sla1, aps1*). These factors are well-established drivers of ubiquitin-dependent cargo sorting and endosomal membrane remodeling processes (101-105).

In contrast, Top2-hc-specific hits emphasized vacuolar lumenal pH control and vesicle tethering/fusion machinery. These included the endosomal $Na^+/H^+$ exchanger (*nhx1*) (106) and V-ATPase V0 subunits (*vph1, vma16, vma11*) that establish appropriate endosomal and vacuolar pH gradients (79). CORVET/HOPS core tethering factors that promote early-to-late endosome and vacuole fusion events were also Top2-hc-specific (*pep3, pep5*) (107). Additional vesicle-traffic mutants included the endosomal SNARE *syn8* (108), the exocytic v-SNARE *snc1* (which recycles via endocytosis), and the endocytic internalization factor *end3* (109). The Top2-hc screen also identified mutants affecting subunits of the Sec62/Sec63 post-translational endoplasmic reticulum translocon (*sec66, sec72*) (**Dataset S3**), which mediate protein translocation into the endoplasmic reticulum (110).

Common hits between the two screens pointed to a shared sorting backbone, including retromer and ESCRT effectors (*vps35, vps24, vps36, vps8*) along with deubiquitination, lipid-handling, and transport factors (*doa4, vps3, cdc50, psd2, rav2, tda3*). This pattern is consistent with shared requirements for cargo deubiquitylation, lipid transfer at membrane contact sites, flippase-assisted trafficking, and V-ATPase assembly (111-114).

**Analysis of TDA genes.** The original Top1-TA SPA screen identified eleven genes of previously unknown function, designated as Topoisomerase I damage affected (TDA) genes (18). In spot tests, most of these *tda* mutants did not exhibit sensitivity to Top2-hc expression alone (***SI Appendix*, Fig. S3*H*, panel i**). However, *tda1* and *tda8* mutants did display sensitivity when challenged by Top2-hc expression together with mAMSA treatment (***SI Appendix*, Fig. S3*H*, panel ii**). The function of Tda8 remains unknown. Tda1 is a protein kinase that is a suppressor of glucose starvation signaling (115).

### Supplemental Methods

#### Plasmid construction

To generate plasmid pSK418 (pRS415-$P_{CUP1}$), which contains the copper-inducible promoter *CUP1* ($P_{CUP1}$), a XhoI-BamHI fragment from pCAS49 (provided by M. Ptashne, MSK) was cloned into the same sites of the *ARS, CEN*, *LEU2* pRS415 plasmid (116) (Stratagene). To generate plasmid pSK420 (pRS415-$P_{CUP1}$-*TOP2*), a PCR-obtained fragment containing yeast *TOP2* from the BY4741 background was cloned into the BamHI and XbaI sites of pSK418. A BamHI-SacI fragment from plasmid p$P_{DED1}$-*top2-F1025Y-R1128G* (19) containing the full coding sequence for *TOP2-hc* was cloned into the same sites of pSK420, replacing *TOP2* and generating pSK778. To generate C-terminally 3×FLAG-tagged Top2 versions, the 3' region of *TOP2* from pSK420 was PCR-amplified, the product digested with NheI-SacII, and ligated into the same sites of pSK420, resulting in pSK814 (pRS415-$P_{CUP1}$-*TOP2-3×FLAG*). The same protocol was followed for tagging *TOP2-hc*, forming pSK815 (pRS415-$P_{CUP1}$-*TOP2-hc-3×FLAG*).

For the yeast selective ploidy ablation (SPA) screen, yeast *TOP2* constructs were inserted into the centromeric vector pRS415 under the control of the copper-inducible *CUP1* promoter. Three related plasmids were used: the empty vector control (pSK418; pRS415-*pCUP1*), a construct expressing either wild-type *yTOP2* (pSK814) or *TOP2-hc* (pSK815), both carrying the same C-terminal 3×FLAG tag. Additional derivatives were created by inserting *GFP* in-frame downstream of the C-terminal 3×FLAG tag, resulting in *yTOP2-3×FLAG-GFP* (pSK842) and *yTOP2-hc-3×FLAG-GFP* (pSK843) for cytological analyses (**Dataset S6**).

For *top2-4* complementation assays, *TOP2* coding sequences were cloned into centromeric pRS316 plasmids under the control of the *CUP1* promoter and followed by the *ADH1* terminator, with the endogenous *URA3* cassette replaced by the *hphMX6* selectable marker. Plasmids in this series included *yTOP2* (pSK1276) or y*TOP2-hc* (pSK1277), as well as the human *TOP2A* WT (pSK1278) and mutants (pSK1279–1281), or *hTOP2B* WT (pSK1282) and mutants (pSK1283–1285) (**Dataset S6**). All human TOP2B constructs used in this study correspond to UniProt splicing isoform Q02880-2, which lacks residues 24–28 relative to the canonical TOP2B sequence. For consistency with other publications, all residue numbering referenced is according to the canonical sequence (UniProt Q02880-1).

The pcDNA5/FRT/TO/CAT vector was used as a platform for cloning cDNAs encoding human TOP2A (pSK955) or TOP2B (pSK957), fused to a C-terminal 2×FLAG tag. The hypercleavage mutations TOP2A-K1140G (pSK956) or TOP2B-K1158G (pSK959) were introduced using the PCR-based Agilent QuikChange Lightning Site-Directed Mutagenesis Kit and verified by sequencing. These constructs allow single-copy Flp recombinase-mediated integration at an FRT site and doxycycline-inducible expression in Flp-In T-REx cells (**Dataset S6**).

For generation of the knock-in mouse targeting constructs, the plasmid cTGM-shRenilla (provided by D. Alonso-Curbelo, S. Lowe laboratory, MSK), was used as the starting backbone and modified to generate the cloning platform cTGM-LSL-IRES2-EGFP (pSK1131). Mouse *Top2a* (pSK1132) and *Top2b* (pSK1134) cDNAs, C-terminally 2×FLAG tagged, were then cloned. The hypercleavage mutations *Top2a-K1137G* (pSK1133) or *Top2b-K1146G* (pSK1135) were introduced by QuikChange and verified by sequencing (**Dataset S6**).

The lentiviral vector pLenti CMV/TO Puro DEST (670-1) (gift from E. Campeau and P. Kaufman; RRID:Addgene_17293) was used to generate constructs expressing N-terminally 3×FLAG-tagged *hTOP2A* or *hTOP2B* alleles in *rtTA*-expressing HeLa and RPE-1 cells. Each plasmid additionally contained an *EF1αcore-EGFP-NLS-P2A-puro* cassette for constitutive *EGFP-NLS* expression and puromycin selection. The resulting vectors encoded *3×FLAG-*

*hTOP2A* (pSK1530), *3×FLAG-hTOP2A-K1140G* (pSK1531), *3×FLAG-hTOP2B* (pSK1532), and *3×FLAG-hTOP2B-K1158G* (pSK1533) (**Dataset S6**).

**Yeast methods**

***Immunoblotting.*** Yeast cultures were centrifuged, supernatant discarded, washed in cold Mili-Q water, fixed with 20% TCA, and stored at −80 °C. Samples were thawed on ice, glass beads were added, and cell lysates were prepared using a FastPrep-24™ classic bead beating grinder and lysis system (MP Biomedicals), 3 cycles at speed 6.0 m/s for 30 s. The lysates were recovered by punching a hole in the bottom of each tube, placing the perforated tube into a new tube, and centrifuged at 1000 × g for 3 min, then the supernatant was discarded. Pellets were thoroughly resuspended in 2× Laemmli buffer and neutralized with 2 M Tris base. After boiling for 5 min, samples were centrifuged at 16,100 × g for 5 min and the supernatant containing the protein extracts passed to a new tube. 10–20 µL were loaded onto 3–8% NuPAGE Tris-Acetate Gels (Thermo Fisher Scientific), run with NuPAGE Tris-Acetate SDS Running Buffer, transferred to PVDF membranes, blocked in 5% non-fat milk PBS-0.1% Tween-20 and incubated with the appropriate primary and secondary antibodies (**Dataset S8**). The Precision Plus Protein Dual Color Standards were used for molecular weight estimation (Bio-Rad). The chemiluminescence signal was captured using ChemiDoc XRS+ and MP imaging systems (Bio-Rad) and ImageLab software (Bio-Rad; version 6.1.0 build 7).

***Cytology.*** Yeast cell images were captured with a Zeiss Axio Observer Z1 Marianas Workstation using a 100×/1.4 NA oil-immersion objective. Rad52-EGFP was imaged in live cells. For Top2-GFP and Top2-hc-GFP visualization, cells were subjected to mild glutaraldehyde fixation, which preserved endogenous GFP fluorescence, and nuclei were counterstained with DAPI. DIC, GFP, and DAPI images were acquired with SlideBook 5.0 (Intelligent Imaging Innovations) and analyzed with Fiji (117).

***Top2cc isolation by ICE assay.*** The ICE assay was conducted as described (52). In brief, yeast cell lysates were prepared with a chaotropic agent lysis buffer in a FastPrep-24™ (MP Biomedicals). Then, they were applied on top of a 150% (w/v) CsCl solution, tubes were sealed, placed in a TN-1865 Neo Angle Rotor (Thermo Scientific), and ultracentrifuged at 42,000 rpm (~157,000 × g) using a Sorvall wX+ Ultra Series ultracentrifuge (Thermo Scientific), for 20 h at 24 °C. The resulting genomic DNA pellets with covalently bound proteins were washed with 70% ethanol and dissolved in 1× TE buffer (10 mM Tris-HCl pH 8.0, 0.1 mM EDTA) for 2 h at room temperature and were quantified using the Qubit dsDNA broad-range assay kit with a Qubit 3.0 fluorometer (Thermo Fisher Scientific). 1 µg of double-stranded DNA was mixed with 25 mM sodium phosphate pH 6.5 buffer, then applied to a 0.45 µm nitrocellulose membrane (Bio-Rad) using a slot-blot vacuum manifold (Bio-Rad) and immunodetected with anti-FLAG (plasmid Top2 or Top2-hc), anti-Top2 (total) and anti-double-stranded DNA (as loading control), followed by incubation with ECL Prime western blotting detection reagent (Amersham) and detection using a ChemiDoc XRS+ imaging system (Bio-Rad).

***Top2-hc and mAMSA sensitivity assays (spot tests).*** Exponentially growing yeast cells in selective medium lacking leucine (SC-LEU) for selection to maintain the plasmids were serially diluted in autoclaved Milli-Q water. The first dilution was adjusted to $OD_{600nm}$ = 0.2, followed by five consecutive 1:5 dilutions. 9 µL of each dilution were spotted onto SC-LEU plates supplemented with 100 µM $CuSO_4$ and/or with mAMSA (MedChemExpress) at indicated concentrations. Plates were incubated at 30 ºC for 3 days and scanned at 24, 40, 44, 48, and 68 h using an EPSON scanner. Images shown correspond to the 44 h time point, which was selected as the most representative. Interspecific cross-complementation assays were performed similarly in the temperature-sensitive *top2-4* background, using hygromycin for selection and plates were incubated at 28 °C and 30 °C (permissive) or 34 °C and 37 °C (non-permissive).

***Selective ploidy ablation (SPA) screen.*** In this method, a plasmid is first introduced into a universal donor strain in which every chromosome carries *URA3* and a galactose-inducible promoter adjacent to its centromere (118). Mating the donor strain in arrayed format to the collection of deletion strains generates a panel of intermediate diploids, which are then serially transferred to plates containing galactose to destabilize the donor chromosomes by transcriptional interference with centromere function, then to plates containing 5-fluoroorotic acid (5-FOA) to select for cells that had lost all of the donor chromosomes (118)

Yeast crosses for the SPA screen were performed by high-density pinning using a Singer RoToR workstation (Singer Instrument Co., Ltd.). Disposable replica pins (RePads) were obtained from Singer, and rectangular Petri plates were obtained from Singer and iGene Supplies, LLC. Screen plate images were captured using a ScanMaker 9800XL flatbed scanner (MicroTek) equipped with the TMA1600 transparent media adapter. Scans were performed using a custom mask with openings for nine rectangular plates to eliminate reflection anomalies (18).

The primary screen contained 4565 open reading frames, 4 colony replicas, 3 plasmids, and 2 conditions (±100 μM $CuSO_4$). These mutants were pin transferred in quadruplicate onto new plates (12.5 × 8.5 cm), creating 32 × 48 grids (1536 colonies total). The SPA protocol was then applied and the resulting haploid strains containing the plasmids were pinned onto selective synthetic medium plates. Therefore, each mutant was represented by four colonies on three different plates (one plate per plasmid) under two copper conditions (SC-LEU or SC-LEU + 100 μM $CuSO_4$), requiring a total of six plates per set of 384 mutants tested (example plates in ***SI Appendix*, Fig. 2*B***). Plate images were scrutinized with the image-analysis software suite *ScreenMill* to get colony size measurements and calculate descriptive statistics for each condition (119). Median-normalized growth values were used to calculate $\log_2$ growth ratios between experimental and vector control colonies (***SI Appendix*, Fig. 2*A* and Dataset S1**). These data approximate a normal distribution, so p-values and z-scores were calculated for each mutant and condition.

To confirm the phenotypes and reduce variability, mutants with $|z| > 2$, plus a few others (technical issues, synthetic lethality phenotypes), up to a total of 523 unique ORFs (some ORFs were repeated), were selected for an additional round of screening. This validation screen contained 546 mutants, 16 colony replicas, 3 plasmids, 2 conditions (±100 μM $CuSO_4$). Representative mutants are shown in ***SI Appendix*, Fig. 2*D*** and full dataset is in **Dataset S1**.

Hits were selected using the more robust output from the validation screen: 110 deletion mutants were hypersensitive to Top2-hc expression ($\log_2$ growth ratio $< -0.5$, $p < 0.05$) and 28 suppressor mutants surprisingly grew better in the presence of Top2-hc as compared to the empty vector control ($\log_2$ growth ratio $> 0.5$, $p < 0.05$).

***Protein structure visualization.*** The yeast Top2-DNA-AMPPNP complex (PDB 4GFH) (120) was used to model the Top2-hc amino acid substitutions using UCSF ChimeraX. For cross-species comparison of the C-gate region, the previous yeast Top2 structure was aligned with human TOP2B in complex with DNA and etoposide (PDB 7YQ8) (121) and human TOP2A DNA-binding/cleavage domain in state 1 (PDB 6ZY5) (122). Figures were prepared in UCSF ChimeraX.

**Mammalian methods**

***Human cell lines.*** HeLa Flp-In™ T-REx™ cells (Thermo Fisher Scientific) were cultured in DMEM high glucose (DMEM-HG) supplemented with 10% FBS and 1% each penicillin and streptomycin, at 37 °C in 5% $CO_2$. To generate isogenic DOX-inducible lines, cells were co-transfected with the pOG44 Flp recombinase expression plasmid and pcDNA5/FRT/TO vectors encoding 2×FLAG-tagged TOP2A or TOP2B constructs (wild-type or hypercleavage mutants). Hygromycin-resistant populations were expanded and verified for DOX-dependent expression of TOP2–2×FLAG proteins following treatment with 1 µg/mL DOX (12–24 h), as evaluated by anti-FLAG immunoblotting.

RPE-1 lines were maintained as for HeLa except using DMEM/F12. A validated *TP53*-knockout RPE-1 cell line was generously provided by P. Jallepalli's laboratory (MSK). To implement a DOX-inducible expression system, RPE-1 *TP53*-KO cells were transduced with a lentiviral vector encoding a constitutively expressed Tet-On 3G transactivator. Lentiviruses were produced in HEK293T cells by cotransfection of the transfer vector with third-generation packaging plasmids. Viral supernatants were collected 48 h post-transfection, filtered (0.45 µm), and applied directly to target cells in the presence of 10 µg/mL polybrene. Tet-On 3G-positive RPE-1 and HeLa Flp-In™ T-REx™ were then transduced with lentiviral vectors expressing CMV-tet$O_2$–regulated 3×FLAG-tagged TOP2A or TOP2B (wild-type or hypercleavage mutants), followed by an IRES–EGFP fluorescent reporter cassette. Puromycin-resistant populations were expanded and verified for DOX-dependent expression of 3×FLAG–TOP2 proteins following treatment with 1 µg/mL DOX (12–24 h), as evaluated by anti-FLAG immunoblotting.

***shRNA screen.*** We employed verified stable HeLa Flp-In™ T-REx™ cell lines with DOX-inducible expression of human *TOP2A-2×FLAG* (wild type (WT) and K1140G) and *TOP2B-2×FLAG* (WT and K1158G), separately. These cells were transduced with a pooled shRNA library targeting the human DNA damage response (hDDR), prepared by the MSK iRNA Core Facility. Pilot transductions were performed to determine the viral titer required to achieve a multiplicity of infection (MOI) of approximately 0.2 (20% transduction efficiency). After three days of selection in 1 µg/mL puromycin, the first time point (T0) was collected, and the cells started to be grown in complete DMEM-HG containing 2 µg/mL DOX and 50 nM etoposide. These conditions were maintained for 20 days, with time points collected and cells passaged every four days. Frozen cell pellets were kept at –80°C for later batch processing.

Genomic DNA was PCR-amplified to recover integrated shRNA cassettes, using vector-specific primers, for T0 and the final time point (TF). The resulting amplified libraries were indexed, pooled, and sequenced on an Illumina HiSeq platform to quantify shRNA representation. QC analysis across the six hDDR sublibraries (hDDR1–6) demonstrated high fidelity, with an average Sanger validation rate of 81.7% and library quality of 84.5%. Importantly, >99% of designed shRNAs were retained in the libraries, with low levels of underrepresentation (mean 1.08%) and negligible overrepresentation (mean 0.06%), ensuring robust coverage for functional screening. Barcode-demultiplexed shRNA counts were assembled into a gene × sample matrix for the baseline (T0) and post-selection time points (TF) in four series: TOP2A, TOP2A-hc, TOP2B and TOP2B-hc. Counts were normalized for library size (CPM) and stabilized with a small pseudocount ($\varepsilon$=1). For each gene and series, we computed $\log_2$ fold-change relative to T0 and summarized multiple hairpins per gene by the median to obtain one gene-level value per sample. Within each subpool × condition (hc or WT) × time (TF), gene-level values were transformed to robust z-scores using the panel's median and median absolute deviation (MAD). To harmonize series that share internal controls, we centered each subpool×condition×time panel on the median z of firefly luciferase and Renilla luciferase negative control hairpins (z_c). This step aligns panels to the same reference without altering the dispersion or relative gene ranking within a panel. Because subpools can carry offsets, we removed subpool-specific biases while preserving any global hc–WT difference. For each isoform and time point separately, we fit a fixed-effects model on a neutral training set that excluded control hairpins and trimmed extremes. We then subtracted the estimated subpool coefficients from all genes in that block (z_c_pfprot = pool-free, condition-protected). Thus, only subpool offsets are removed; the explicit condition term prevents over-correction of true hc–WT signal. Unless stated otherwise, all figures and statistics use z_c_pfprot. Anchored and corrected z distributions were centered near zero with comparable spread across series, as assessed by density overlays and summary statistics. Controls were rarely called at $z \leq -1.85$ (false-positive rate $\lesssim 1\%$ across series), supporting specificity. Results were stable at nearby thresholds (−1.75 more permissive; −2.0 stricter); −1.85 provided the best balance between sensitivity to strong depletions and control specificity and was adopted as the final cutoff. The pipeline operates at the gene level (median across hairpins), is panel-local (per subpool×condition×time), and relies only on shared controls for anchoring, providing portability across libraries and screens. Analyses were performed in R (v4.3.1, R core team 2023) using dplyr, tidyr, stringr, and readr (tidyverse) (123), ggplot2 with ggrepel for visualization, and base stats for modeling and z-scoring.

***TOP2cc isolation by DUST assay.*** Detection of ubiquitylated and SUMOylated TOP2 cleavage complexes (DUST) assay was performed as described (61) with modifications. Briefly, $0.5 \times 10^6$ cells were seeded in 10-cm plates and treated the following day with 1 µg/mL DOX for 24 h. On the second day, cells were incubated in fresh DMEM-HG containing 1 µg/mL DOX and the proteasome inhibitor bortezomib (10 µM) for 1 h, followed by an additional 1 h co-treatment with the indicated concentration of etoposide or vehicle (DMSO). Cells were rapidly washed with ice-cold PBS and lysed directly in 1 mL of DNAzol or TRIzol. Lysates were collected by scraping, transferred to a microcentrifuge tube and homogenized by repeated pipetting. Ice-cold 100% ethanol was added, samples were thoroughly mixed, and nucleic acids were precipitated by centrifugation (16,100 × g, 15 min, 4 °C). Pellets were washed three times with 75% ethanol, briefly air-dried, and resuspended in 500 µL 1× TE buffer with gentle rotation overnight at 4 °C.

Samples were sonicated (25% amplitude, 10 s on/10 s off pulses, 3 cycles), clarified by centrifugation (16,100 × g, 5 min, 4 °C), and treated with RNase A (100 µg/mL, 30 min, room temperature). DNA was re-precipitated by addition of sodium acetate (0.3 M final) and ethanol (2–2.5 volumes), incubated at –20 °C overnight, centrifuged (16,100 × g, 15 min, 4 °C), washed with 70% ethanol, and resuspended in 1× TE buffer. DNA concentration was determined using Qubit dsDNA broad-range assays. Samples were supplemented with $MgCl_2$ and digested with Turbonuclease (1–2 h, room temperature). LDS sample buffer containing freshly added DTT was then added, and samples were heated at 95 °C for 5 min, followed by centrifugation (16,100 × g, 5 min). Supernatants were transferred to fresh tubes.

Sample volumes corresponding to 1–2.5 µg DNA were resolved by SDS–PAGE on NuPAGE 3–8% Tris-acetate gels using NuPAGE Tris-Acetate SDS running buffer (Thermo Fisher Scientific), transferred to PVDF membranes and analyzed by immunoblotting. Membranes were blocked in 5% non-fat milk in PBS containing 0.1% Tween-20 and incubated with HRP-conjugated anti-FLAG or anti-TOP2 antibodies (**Dataset S8**). Detection was performed using ECL Prime (Amersham), and chemiluminescence was acquired on a ChemiDoc MP imaging system (Bio-Rad) and ImageLab software (Bio-Rad; version 6.1.0 build 7).

***Cell viability assays.*** HeLa, RPE-1, or MEFs were seeded at 5,000 cells per well in 96-well plates and allowed to adhere overnight. The following day, 1 µg/mL DOX was added where indicated to induce TOP2 construct expression, and cells were treated with vehicle (DMSO) or increasing concentrations of etoposide (0.25, 0.5, 1, 2.5, 5, 10, or 50 µM) for 72 h in the presence or absence of DOX. Each condition was performed in three technical replicates per cell line. At the end of treatment, medium was replaced with CellTiter-Blue reagent (Promega) prepared according to the manufacturer's instructions, and fluorescence was measured after 3–8 h of incubation using a SpectraMax iD5 microplate reader (Molecular Devices). Viability values were normalized to untreated wild-type controls within each experiment. Data were analyzed and plotted in GraphPad Prism (v10–11). Individual replicate values are displayed, and statistical comparisons between genotypes at each dose were performed using two-tailed Welch's t-tests.

***Immunoblotting.*** Adherent mammalian cells were trypsinized, neutralized in cold complete medium, collected by centrifugation, washed in cold PBS without $Ca^{2+}$/$Mg^{2+}$, resuspended in cold PBS without $Ca^{2+}$/$Mg^{2+}$ supplemented with Halt protease inhibitors, snap-frozen on dry ice, and stored at −80 °C until processing. After partial thawing, 2× SDS sample buffer [100 mM Tris-HCl (pH 6.8), 4% SDS, 20% glycerol, 100 mM DTT, and bromophenol blue] was added to a final 1× concentration. For DNA damage-response immunoblots, cells were washed in cold PBS without $Ca^{2+}$/$Mg^{2+}$, lysed directly with 2× sample buffer and collected by scraping. Mouse organs were dissected, washed in PBS, snap-frozen on dry ice, and stored at −80 °C until processing. Frozen tissue was cut into ~30-mg pieces and transferred to microcentrifuge tubes; for some organs, tissue was first finely chopped with a blade to facilitate subsequent homogenization with a plastic pestle. Tissue samples were lysed directly in 600 µL of 2× SDS sample buffer and mechanically homogenized until well disrupted.

All lysates were sonicated briefly (20% output, 10-s on/10-s off pulses, 2 cycles), heated at 95 °C for 5 min for cell lysates or at 70 °C for 10 min for tissue lysates, and then clarified by centrifugation at 16,100 × g for 5 min. Protein concentration was measured using the RC DC Protein Assay (Bio-Rad) according to the manufacturer's instructions, and equivalent amounts of protein (10–30 µg) were loaded per lane. Samples were resolved by SDS-PAGE on NuPAGE 4–12% Bis-Tris gels run with NuPAGE MES or MOPS SDS Running Buffer or on 3–8% Tris-acetate gels run with NuPAGE Tris-Acetate SDS Running Buffer (Thermo Fisher Scientific), transferred to PVDF membranes, and analyzed by immunoblotting. Membranes were blocked in 5% non-fat milk or BSA in PBS or TBS-0.1% Tween-20 and incubated with the primary antibody overnight at 4 °C, washed, incubated with HRP-conjugated secondary antibodies (**Dataset S8**), washed again and developed with ECL Prime (Amersham). Chemiluminescence was captured using a ChemiDoc MP imaging system (Bio-Rad) and ImageLab software (Bio-Rad; version 6.1.0 build 7).

***Immunofluorescence staining.*** MEFs were seeded, cultured and treated in glass-bottom, black-wall 24-well plates and washed with PBS prior to fixation in 4% paraformaldehyde (PFA) in PBS for 10–15 min at room temperature. After three washes with cold PBS, cells were permeabilized with 0.25% Triton X-100 in PBS for 10 min at room temperature and washed three times in PBS. Samples were blocked in 5% BSA in PBS-0.1% Tween-20 (PBS-T) and incubated with primary antibodies diluted in 1% BSA PBS-T overnight at 4 °C (**Dataset S8**). After three PBS-T washes, cells were incubated with fluorophore-conjugated secondary antibodies (Alexa Fluor; Invitrogen) diluted in 1% BSA in PBS-T for 1 h at room temperature, protected from light. Wells were washed three times in PBS-T, after which mounting medium containing DAPI (Vectashield) was added directly to each well, and were covered with 15-mm circular glass coverslips. Samples were imaged on a Zeiss Axio Observer Z1 Marianas Workstation using 63×/1.4 NA and 100×/1.4 NA oil-immersion objectives. Image acquisition was performed using SlideBook 6.0 and 2025 (Intelligent Imaging Innovations), and images were analyzed with Fiji (117).

***Mice.*** Mice were housed in solid-bottom, polysulfone, individually ventilated cages (IVCs) (Thoren Caging Systems, Hazelton, PA) on autoclaved aspen-chip bedding (PWI Industries Canada, Quebec, Canada); γ-irradiated feed (LabDiet 50531, PMI, St Louis, MO) and acidified reverse osmosis water (pH 2.5 to 2.8) were provided ad libitum. The cages also contained Nestlets®, EnviroDri®, and/or EnviroPaks® as environmental enrichment. The IVC system was ventilated at approximately 30 air changes hourly. HEPA-filtered room air was supplied to each cage and the rack effluent was exhausted directly into the building's exhaust system. Cages were changed weekly in either a HEPA-filtered vertical flow change station or a Class 2 Type A biological safety cabinet. The animal holding room was maintained at 21.5 ± 1 °C, relative humidity between 30% and 70%, and a 12:12 h light:dark photoperiod.

CRE-conditional, DOX-inducible *Top2-hc knock-in* mouse models were generated using FlpE-mediated recombinase-mediated cassette exchange in mouse embryonic stem cells (mESCs). cDNAs encoding mTOP2A-hc-2×FLAG (pSK1133) or mTOP2B-hc-2×FLAG (pSK1135) were cloned into the *Col1a1* targeting vector (cTGM) and electroporated into KH2 mESCs to achieve single-copy transgene insertion downstream of the endogenous *Col1a1* locus (54, 55). Hygromycin-resistant clones were isolated, and three independent lines per construct were submitted for karyotype analysis (Molecular Cytogenetics Core Facility, MSK). Clones with normal male karyotypes were selected for blastocyst injection. Targeted mESCs were microinjected into blastocysts derived from NCI C57BL/6-cBrd/cBrd/Cr (C57BL/6 albino) mice and implanted into CD-1 pseudopregnant females, resulting in chimeric offspring (Rodent Genetic Engineering Laboratory, NYU Langone). High-percentage chimeras were bred to C57BL/6J mice (The Jackson Laboratory) to establish germline transmission. Founder animals carrying confirmed *mTop2a-hc-2×FLAG* or *mTop2b-hc-2×FLAG* alleles were crossed with *Rosa26-CAGs-LSL-rtTA3-IRES2-mKate2* (RIK; JAX #029633) mice (40) to enable DOX-dependent expression. Heterozygous mice were backcrossed to the C57BL/6J background for at least ten additional generations. To remove loxP-flanked STOP (LSL) cassettes *in vivo*, *Col1a1-TRE-LSL-Top2a/b-hc-2×FLA*G-IRES2-EGFP *Rosa26-CAGs-LSL-RIK* mice were crossed with the germline Cre-deleter strain *E2a-Cre* (JAX #003724) (124). This strategy yielded animals with constitutive, widespread expression of rtTA3 and mKate2, permitting DOX-inducible expression of TOP2A/B-hc-2×FLAG and EGFP. For induction experiments, mice were switched to a DOX-containing chow (TD.01306, 625 mg/kg, ENVIGO and Bio-Serv) for 5–7 days or until reaching a humane endpoint of 20% weight loss (whichever occurred first). Animals were euthanized by carbon dioxide overdose, followed by tissue collection.

***Isolation and Culture of MEFs.*** Primary MEFs were isolated from E13.5 embryos. Pregnant females were euthanized at 13.5 days post coitum, and uterine horns were dissected and transferred to PBS without $Ca^{2+}/Mg^{2+}$. Individual embryos were isolated, and placentas, fetal liver (retained for genotyping), head, and internal organs were removed. Embryonic tissue was minced and dissociated in 0.05% trypsin–EDTA for 15–20 min at 37 °C with intermittent trituration. Trypsin was neutralized with DMEM-HG supplemented with 10% FBS and 1% penicillin–streptomycin. DNase I was added to reduce viscosity. Cells were pelleted (300 × g, 5 min), resuspended in DMEM-HG, and plated onto 10-cm dishes (one embryo per dish). Cultures were maintained at 37 °C with 5% $CO_2$, and medium was replaced after 24 h to remove debris. Primary MEFs were expanded and used at early passage (P1–P2) or cryopreserved. For immortalization, early-passage MEFs were transduced with lentiviral vectors encoding SV40 large T antigen (SV40T) and selected to establish stable immortalized lines. Immortalized MEFs were maintained under the same culture conditions as primary cells and used for subsequent experiments.

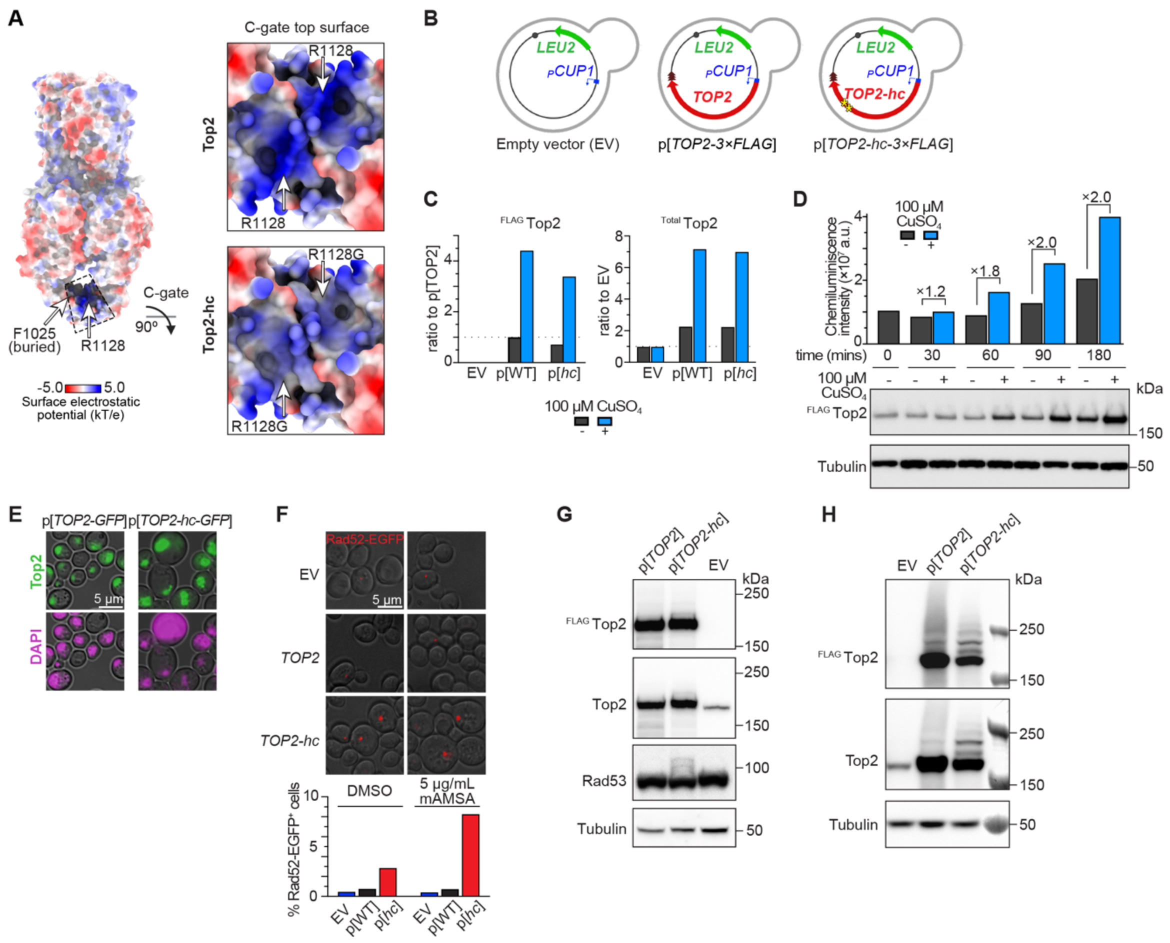


***SI Appendix*, Fig. S1: Validation of the yeast *TOP2-hc* allele.**

(A) Reduced positive electrostatic surface potential at the C-gate from Top2-hc substitutions. Left, overall structure of the Top2 homodimer (PDB 4GFH) highlighting the boxed C-gate interface. Right, magnified views of the C-gate dimer interface shown for wild-type Top2 (top) and Top2-hc (bottom). The reduced local positive surface potential may contribute to destabilizing C-gate interactions, altering the cleavage–religation equilibrium, and increasing the persistence of Top2cc.

(B) Plasmid schematics for low-copy ARS–CEN vectors for empty vector control (EV) and plasmids expressing wild-type *TOP2* or *TOP2-hc* with a C-terminal triple 3×FLAG tag under the control of a copper-inducible promoter ($P_{CUP1}$).

(C) Baseline and copper-induced expression of $^{FLAG}$Top2 or $^{FLAG}$Top2-hc. Quantification of immunoblot in **Fig. 1*A*** is shown, normalized to the tubulin loading control and displayed relative to baseline $^{FLAG}$Top2 (left) or relative to total Top2 in the EV control (right).

(D) Time course of copper-induced $^{FLAG}$Top2 expression .

(E) Nuclear localization and cytotoxicity of Top2-hc expression. Representative images are shown of Top2 or Top2-hc with a C-terminal GFP fusion in a wild-type strain after 20 hr of copper induction. Top2-hc overexpression caused cellular morphologies associated with cytotoxicity, such as swelling, lysis, and loss of the rounded shape and the membrane integrity.

(F) Rad52-EGFP foci induction upon Top2-hc expression. These foci mark sites of active homologous recombination-mediated repair (125). Exponentially growing cultures were treated for 5 hr with 100 µM $CuSO_4$ and either 5 µg/mL mAMSA or DMSO. Representative images (top)

and quantification (bottom) are shown. Values indicate EGFP-positive cells/total cells per condition (n = 6/1277, 15/1917, 70/2442, 16/3783, 10/1362, and 148/1787, respectively).
(G) Activation of the DNA damage checkpoint kinase Rad53 by Top2-hc expression. Phosphorylation-dependent Rad53 activation results in accumulation of slower-migrating forms (126). Immunoblots are shown of whole-cell extracts from exponentially growing cultures harvested 4.5 hr after dilution to $OD_{600}$ = 0.2 in medium supplemented with 100 μM $CuSO_4$.
(H) Accumulation of slower-migrating Top2-hc species, consistent with posttranslational modification (likely ubiquitination and/or sumoylation) associated with Top2cc processing (29, 127). Whole-cell extracts were prepared 24 hr after dilution into copper supplementation medium.

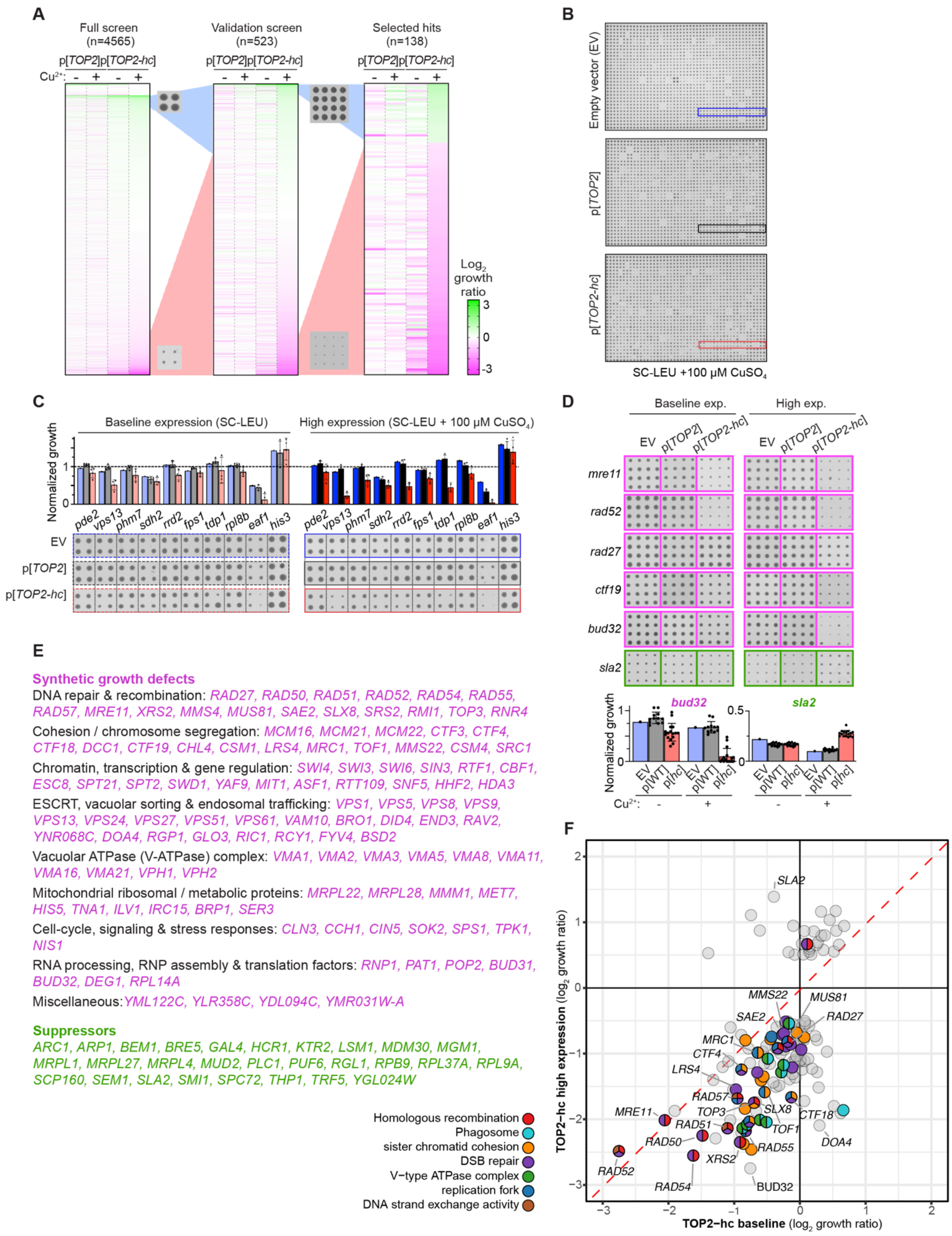
A
Full screen (n=4565)
Validation screen (n=523)
Selected hits (n=138)
p[TOP2] p[TOP2-hc]
Cu2+: - + - +
Log2 growth ratio
3
0
-3
B
Empty vector (EV)
p[TOP2]
p[TOP2-hc]
SC-LEU +100 μM CuSO4
C
Baseline expression (SC-LEU)
High expression (SC-LEU + 100 μM CuSO4)
Normalized growth
pde2 vps13 phm7 sdh2 rrd2 fps1 tdp1 rpl8b eaf1 his3
EV
p[TOP2]
p[TOP2-hc]
D
Baseline exp.
High exp.
EV p[TOP2] p[TOP2-hc]
mre11
rad52
rad27
ctf19
bud32
sla2
Normalized growth
EV p[WT] p[hc]
Cu2+: - +
E
Synthetic growth defects
DNA repair & recombination: RAD27, RAD50, RAD51, RAD52, RAD54, RAD55, RAD57, MRE11, XRS2, MMS4, MUS81, SAE2, SLX8, SRS2, RMI1, TOP3, RNR4
Cohesion / chromosome segregation: MCM16, MCM21, MCM22, CTF3, CTF4, CTF18, DCC1, CTF19, CHL4, CSM1, LRS4, MRC1, TOF1, MMS22, CSM4, SRC1
Chromatin, transcription & gene regulation: SWI4, SWI3, SWI6, SIN3, RTF1, CBF1, ESC8, SPT21, SPT2, SWD1, YAF9, MIT1, ASF1, RTT109, SNF5, HHF2, HDA3
ESCRT, vacuolar sorting & endosomal trafficking: VPS1, VPS5, VPS8, VPS9, VPS13, VPS24, VPS27, VPS51, VPS61, VAM10, BRO1, DID4, END3, RAV2, YNR068C, DOA4, RGP1, GLO3, RIC1, RCY1, FYV4, BSD2
Vacuolar ATPase (V-ATPase) complex: VMA1, VMA2, VMA3, VMA5, VMA8, VMA11, VMA16, VMA21, VPH1, VPH2
Mitochondrial ribosomal / metabolic proteins: MRPL22, MRPL28, MMM1, MET7, HIS5, TNA1, ILV1, IRC15, BRP1, SER3
Cell-cycle, signaling & stress responses: CLN3, CCH1, CIN5, SOK2, SPS1, TPK1, NIS1
RNA processing, RNP assembly & translation factors: RNP1, PAT1, POP2, BUD31, BUD32, DEG1, RPL14A
Miscellaneous: YML122C, YLR358C, YDL094C, YMR031W-A
Suppressors
ARC1, ARP1, BEM1, BRE5, GAL4, HCR1, KTR2, LSM1, MDM30, MGM1, MRPL1, MRPL27, MRPL4, MUD2, PLC1, PUF6, RGL1, RPB9, RPL37A, RPL9A, SCP160, SEM1, SLA2, SMI1, SPC72, THP1, TRF5, YGL024W
Homologous recombination
Phagosome
sister chromatid cohesion
DSB repair
V−type ATPase complex
replication fork
DNA strand exchange activity
F
TOP2−hc high expression (log2 growth ratio)
TOP2−hc baseline (log2 growth ratio)
SLA2 MMS22 MUS81 SAE2 RAD27 MRC1 CTF4 LRS4 RAD57 MRE11 TOP3 SLX8 RAD51 CTF18 TOF1 RAD50 RAD55 DOA4 XRS2 RAD52 RAD54 BUD32

***SI Appendix*, Fig. S2. SPA screen workflow.**
(A) Heatmaps summarizing results from the full screen, validation screen, and final selected hits. Columns show $log_2$ growth ratio scores for p[*TOP2*] and p[*TOP2-hc*] under baseline (-$Cu^{2+}$) and high expression (+$Cu^{2+}$) conditions. Rows are ordered according to p[*TOP2-hc*] scores under high-expression conditions. Color scale indicates $log_2$ growth ratio: magenta denotes synthetic growth defects, white indicates no significant change, and green indicates suppressors.
(B,C) Representative plate images (B) and colony size quantification (C, showing regions highlighted in panel B) from the full screen. In C, the average of four colonies is shown for each strain and condition, normalized to the median colony size of the corresponding plate. Error bars represent SD, and individual dots indicate single-colony measurements.
(D) Examples from the validation screen, showing representative synthetic sick mutants (magenta) and a suppressor mutant (green). Quantification is shown below for *bud32* and *sla2*. Bars represent mean ± SD normalized to the median colony size of the corresponding plate; dots represent individual colonies (n = 16).
(E) Functional categorization of the final 138 selected hits. Synthetic growth-defect hits (magenta) are grouped by annotated biological processes, and suppressors are show in green.
(F) Scatterplot of $log_2$ growth ratios for final selected hits comparing baseline (x axis) versus high-expression (y axis) conditions for p[*TOP2-hc*]. Mutants exhibiting enhanced synthetic sickness under high expression deviate downward from the diagonal. Functional categories are color-coded as indicated.

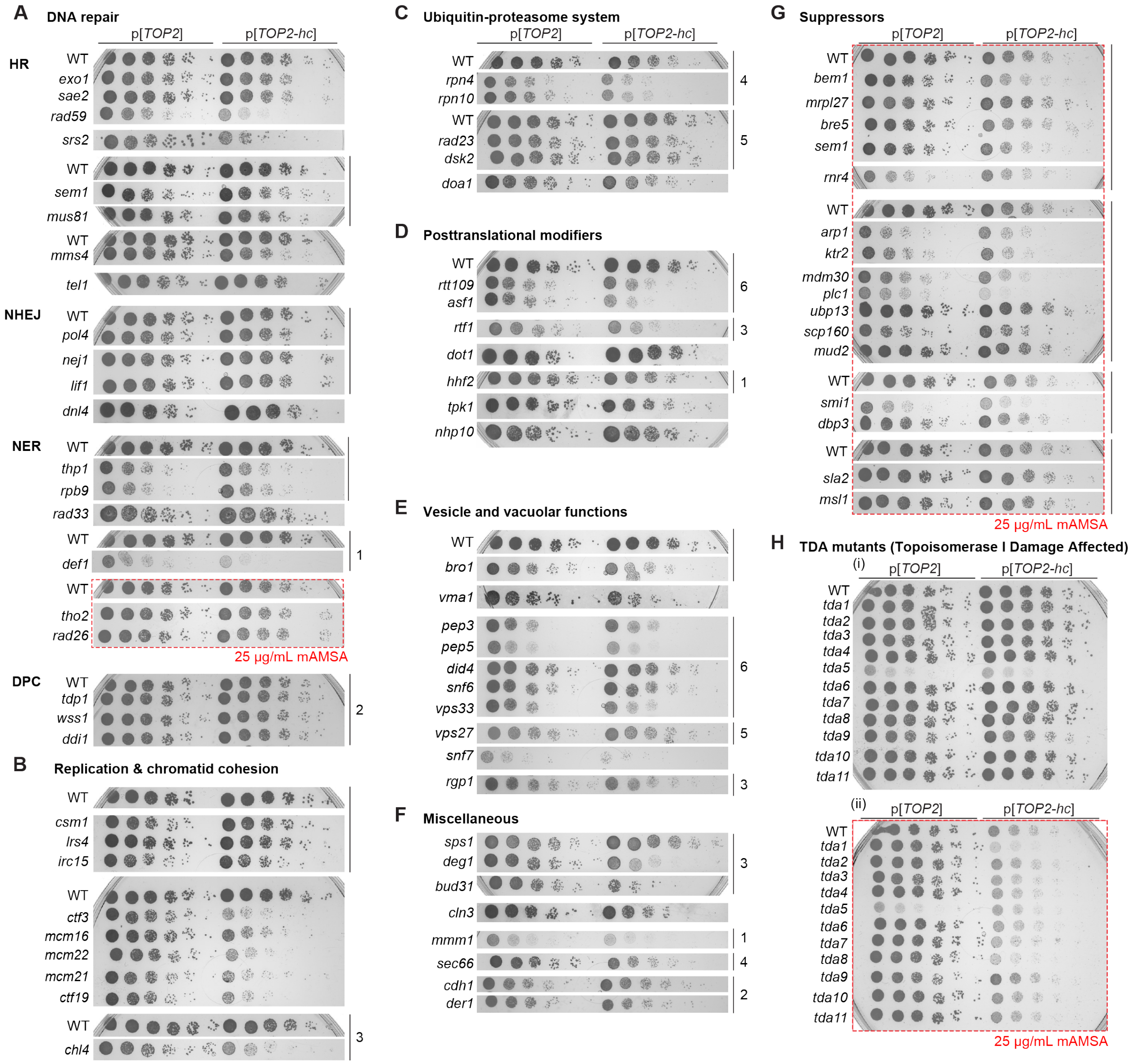


***SI Appendix*, Fig. S3. Spot-test validation of SPA screen results.**

(A-G) Spot-dilution assays as in **Fig. 2C** for mutants identified in the Top2-hc SPA screen, grouped according to functional categories. All plates contained 100 µM $CuSO_4$. Where indicated (red dashed boxes), plates also contained 25 µg/mL mAMSA to assess combinatorial sensitivity. Wherever possible, strains that were contiguous on the original plate are shown together as a continuous image. When relevant strains were not contiguous on the plate, they are shown as separate strips; vertical bars at the right indicate strips derived from the same source plate. Matching numbers to the right identify strips from the same plate that appear in different panels.

(H) Insensitivity of most TDA (Topoisomerase I damage affected) deletion mutants to Top2-hc expression. Spot-dilution assays were conducted in the presence of 100 µM $CuSO_4$ without (i) or with (ii) 25 µg/mL mAMSA.

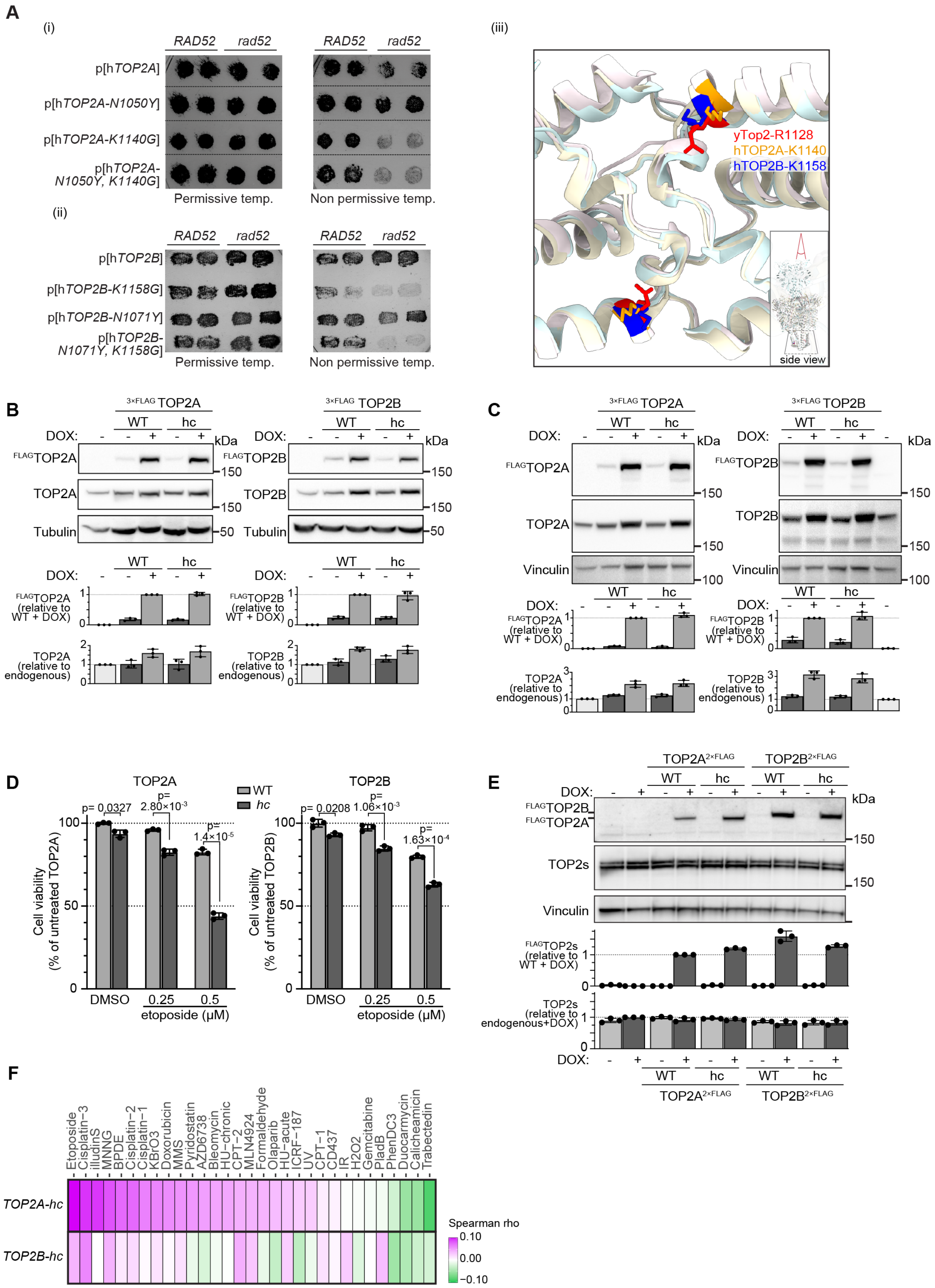


***SI Appendix*, Fig. S4. Verification of human *TOP2* hypercleavage phenotypes.**

(A) Single and double C-gate substitutions in human TOP2A (i) and TOP2B (ii), evaluated by complementation of *top2-4* in *RAD52* and *rad52* yeast strains. Cells were grown for 36 h on selective plates supplemented with 100 µM $CuSO_4$ at permissive (28 ºC) and non-permissive temperatures (35 ºC). C-gate structures are superimposed for the yeast and human proteins (iii), highlighting the conserved residue altered in single-mutant hc variants.
(B) DOX-inducible expression of FLAG-tagged human TOP2A (left) and TOP2B (right) in lentivirally transduced RPE-1 cells. Immunoblots are shown of whole cell extracts prepared after 24 h of DOX exposure (2 µg/mL), probed with anti-FLAG and isoform-specific antibodies. Tubulin serves as a loading control. Quantification of chemiluminescence signal is below (mean ± SD of three replicates). Total TOP2 protein levels were not significantly increased in the absence of DOX relative to control cells lacking expression constructs ($p \geq 0.22$, t tests). DOX induction increased FLAG-TOP2 levels approximately sixfold, with hc variants expressed comparably to WT constructs. Total TOP2 protein increased ~1.6–1.8-fold upon induction, indicating that expression levels were comparable to those of the endogenous proteins.
(C) DOX-inducible expression of FLAG-tagged human TOP2A (left) and TOP2B (right) in lentivirally transduced HeLa cells. Immunoblots are shown as in panel B, except that vinculin serves as the loading control and quantification is expressed relative to the indicated internal controls.
(D) Intrinsic cytotoxicity and enhanced sensitivity to etoposide from TOP2-hc expression in HeLa cells. Cell viability was assessed as in **Fig. 3D** for lentivirally transduced HeLa cells expressing WT or hc TOP2A (left) or TOP2B (right) following DOX induction and treatment with increasing concentrations of etoposide for 3 days. Viability is expressed as percentage of untreated WT controls (with DMSO, without DOX). Bars represent mean ± SD of three replicates; individual data points are shown. P values are from two-tailed Welch's t-tests.
(E) DOX-inducible expression of FLAG-tagged TOP2A and TOP2B in the stable isogenic HeLa Flp-In T-REx cell lines used for shRNA screens. Immunoblots are shown as in panel B, except that vinculin serves as the loading control and total TOP2 levels were assessed with a pan-TOP2 antibody rather than isoform-specific antibodies.
(F) Correlation (Spearman rho) of TOP2A-hc and TOP2B-hc shRNA screen signatures with CRISPR sensitivity profiles for a panel of genotoxic agents (8).

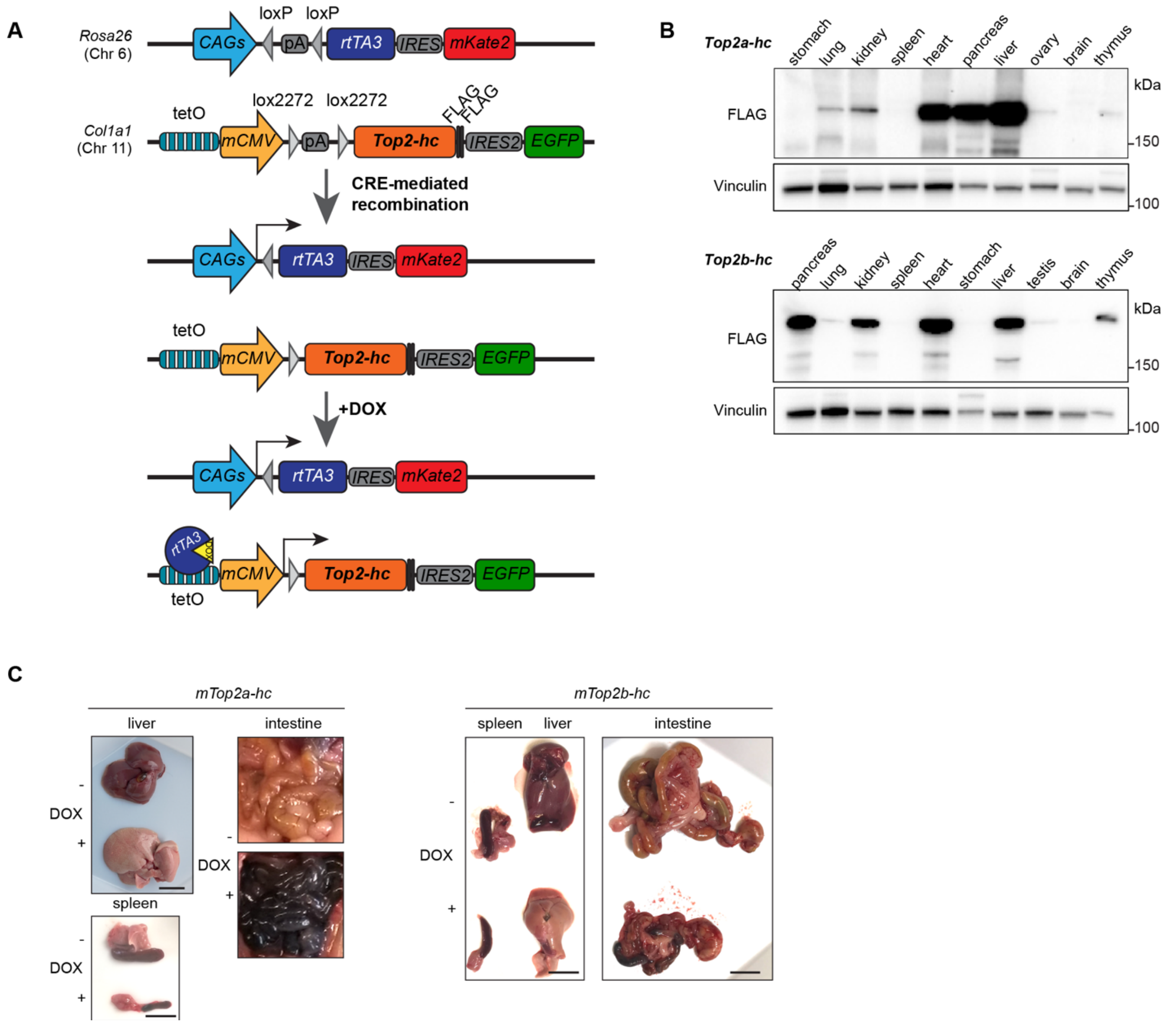


***SI Appendix*, Fig. S5. *Top2-hc* transgenic mouse models.**
(A) Cre-dependent and DOX-inducible *Top2-hc* alleles enable controlled expression *in vivo*. Schematics illustrate the knock-in strategy in which Cre-mediated recombination activates constitutive rtTA3 (and mKate2) expression, permitting DOX-dependent expression of *Top2a-hc* or *Top2b-hc* together with EGFP. pA, transcript cleavage and polyadenylation sequence.
(B) Tissue-variable expression of TOP2A-hc (top) or TOP2B-hc (bottom) after DOX induction. Immunoblots are shown of whole-tissue extracts from DOX-fed mice that carry whole-body excision of the *rtTA3* and *Top2-hc* lox-STOP-lox cassettes. The observed heterogeneity in expression levels is likely attributable to differences in DOX exposure, rtTA/promoter activity, and TOP2 protein stability across tissues. Vinculin serves as a loading control.
(C) Gross pathology representative images of spleen, liver, and intestinal tissues showing gross differences between mice maintained on regular chow (DOX–) or fed DOX (625 mg/kg) for 7 days (DOX+).

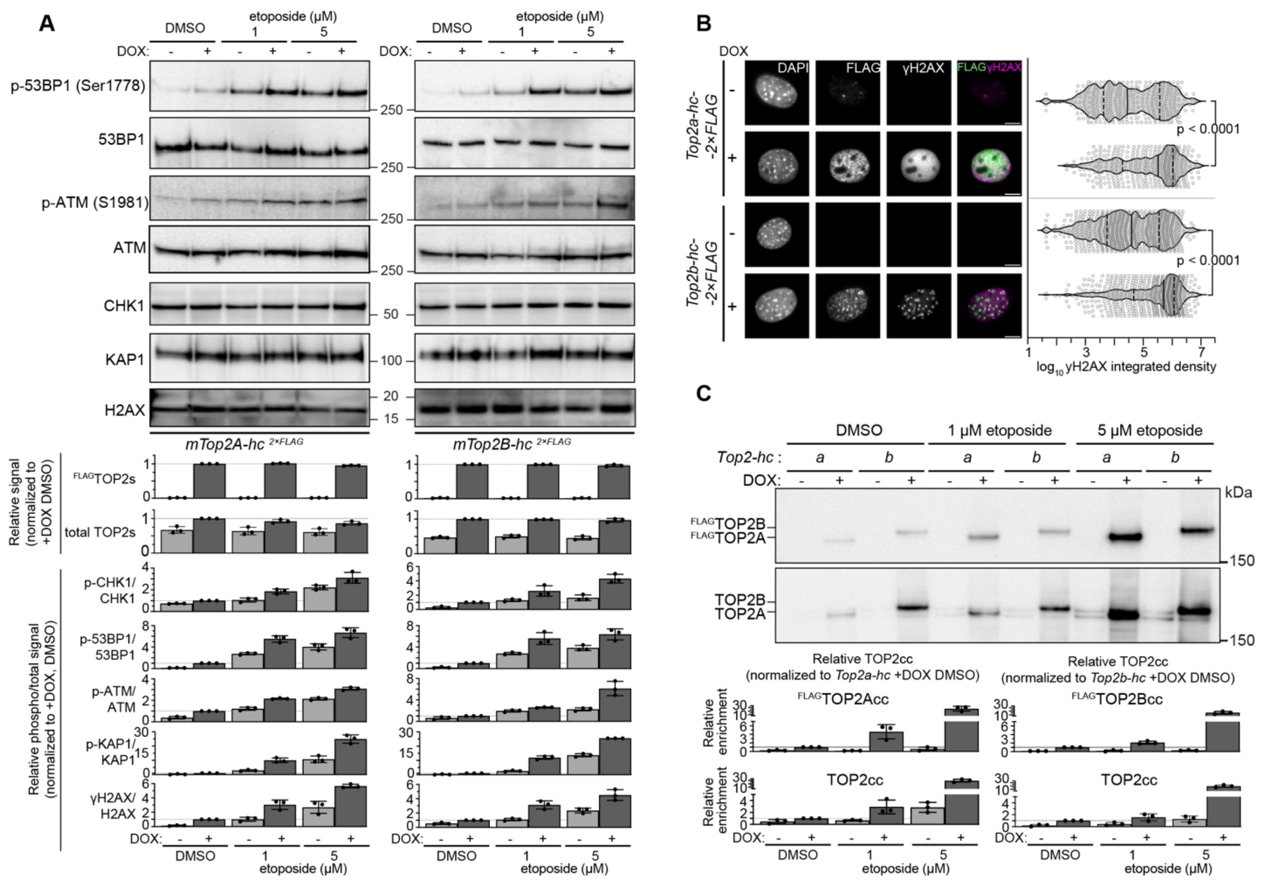


***SI Appendix*, Fig. S6. DNA damage from TOP2-hc expression in immortalized MEFs.**

(A) DNA damage response. Additional immunoblots and quantification (mean ± SD from three replicates) are shown for the experiment in **Fig. 5C**.

(B) γH2AX induction. Representative immunofluorescence micrographs (left) and quantification (beeswarm and violin plots, right) are shown. Each dot represents one cell; solid black bars indicate medians, dashed lines indicate the 25th and 75th percentiles; p values are from two-tailed Mann–Whitney tests).

(C) TOP2cc detection by DUST assay. Cells were treated with or without DOX for 24 h, followed by 2-h treatments with DMSO or etoposide (1 or 5 μM). DUST precipitates were immunoblotted with anti-FLAG or pan-TOP2 antibodies. Quantification below shows mean ± SD of three replicates.
